# Molecular basis of mitochondrial leucine transport by human Sideroflexin 1

**DOI:** 10.64898/2026.06.18.732944

**Authors:** Fiona M. Fitzpatrick, Daniel T. D. Jones, Hannah Aguirre, Benjamin Fry, Elena T. Kämmerer, Sijie Tan, Luise E. Veelken, Wen Han Tong, Samuel Block, S. Ali Aghvami, Edmund R. S. Kunji, Nicholas Polizzi, Stephen C. Blacklow, Nora Kory

## Abstract

Leucine is a central nutrient signal and a ketogenic amino acid that fuels metabolism, yet how it is imported into mitochondria remains incompletely defined. Sideroflexins are conserved inner mitochondrial membrane proteins implicated in amino acid transport, but their mechanism and substrate specificity remain unclear. Using cryogenic electron microscopy, we determined the structure of SFXN1 in its matrix-open conformation. AlphaFold and Boltz co-folding of SFXN1 with a library of human metabolites identified leucine as a candidate substrate, findings supported by thermal stability measurements and mitochondrial transport assays. Comparison with a cytoplasmic-open-model reveals an alternating-access “toggle-switch” mechanism of transport. Together, these findings uncover the molecular basis of leucine transport by SFXN1 and provide a framework for understanding its role in metabolism and disease.

## Introduction

Mitochondria couple oxidative phosphorylation to metabolic homeostasis by controlling the exchange of metabolites across the inner membrane. Because this membrane is impermeable to metabolites and ions, dedicated transport proteins define the chemical composition of the matrix and thereby determine the substrate pools that sustain catabolic and anabolic reactions. Despite their central importance, the molecular basis and substrate specificities of many mitochondrial inner membrane transporters remain unresolved, limiting mechanistic understanding of mitochondrial metabolism and obscuring how defects in these transport processes contribute to disease.

Leucine, an essential amino acid obtained from the diet, is reversibly transaminated to α-ketoisocaproate by branched-chain aminotransferases that exhibit compartment and tissue-specific expression (*1–3*). Mitochondrial leucine transamination is catalysed by BCAT2, the dominant enzyme in humans and the first step in a pathway that generates acetoacetate and acyl-CoA intermediates that sustain tricarboxylic acid cycle flux (*1–5*), particularly in oxidative, mitochondria-rich tissues such as skeletal muscle, heart and brown adipose tissue (*6*). Because these oxidative steps occur mainly in mitochondria, transport of leucine into the mitochondrial matrix constitutes a critical control point in leucine catabolism. In parallel, leucine functions as a nutrient signal that activates mTOR-dependent anabolic programs, placing leucine metabolism at the interface of bioenergetics and growth regulation (*7–9*). Impaired metabolism of branched-chain amino acids, including leucine, underlies maple syrup urine disease (*10*), an inborn error of metabolism. Dysregulated branched-chain amino acid metabolism has been linked to obesity, type 2 diabetes and cancer, in which these amino acids are increasingly utilized to support metabolic demands (*11*, *12*).

The sideroflexins (SFXNs) comprise a conserved family of multi-pass mitochondrial inner membrane proteins, with five paralogues in humans. Genetic studies have linked SFXN family members to iron homeostasis (*13–16*) and mitochondrial dysfunction (*17–19*), and SFXN1 was recently proposed to function as a serine transporter required for one-carbon metabolism, with SFXN3, a closely related paralogue, exhibiting redundant function (*20*). However, the structural basis of SFXN1 function and its substrate preferences remain unclear.

Here we determined the cryogenic electron microscopy (Cryo-EM) structure of human SFXN1 in a matrix-open conformation, revealing a six-transmembrane-helix transporter architecture that encloses a central cavity that extends towards the membrane midplane. Structural bioinformatics using a human metabolite library and SFXN1 co-folding identified hydrophobic amino acids as high-confidence candidate substrates and defined a putative binding mode within the cavity. Consistent with this prediction, these amino acids thermostabilize SFXN1, and reintroduction of SFXN1 into SFXN1/SFXN2 loss-of-function backgrounds increased both the capacity and rate of leucine uptake in isolated mitochondria. Together, these data establish a structural basis for mitochondrial leucine import by SFXN1. This study also provides a blueprint for integrating experimental structure determination with state-of-the-art protein–ligand co-folding to uncover transporter substrates and mechanisms.

### Structure of human Sideroflexin 1

We set out to determine the first structure of a SFXN protein. Despite optimizing heterologous expression of human SFXN1 in *Saccharomyces cerevisiae* to yield highly pure, monomeric protein (fig. S1A), crystallization was not successful. The small size of SFXN1 (32 kDa) also posed challenges for single-particle Cryo-EM analysis.

To facilitate structural determination by Cryo-EM, we engineered apocytochrome b562 (bRIL) – SFXN1 fusion proteins to increase the protein mass of the particles used for analysis (*21*, *22*). We tested insertions of bRIL at the N-terminus or within loops connecting different predicted transmembrane helices (fig. S1B), finding that insertion of bRIL between TM1 and TM2 was well tolerated (fig. S1B). To enhance structural rigidity at the fusion junction, bRIL insertions were designed *in silico* and tested in small-scale to maintain continuous extension of SFXN1 transmembrane helices (see Methods (*23*), fig. S1C and D). We selected three bRIL fusions for large scale purification and complexation with an anti-bRIL Fab and anti-Fab nanobody (*22*, *24–26*). SFXN1-bRIL9 formed a stable complex and was monodisperse during size exclusion chromatography (fig. S1E). Single particle Cryo-EM analysis led to a dataset refined to a global resolution of 3.2 Å (Figs. S2 and S3), with high resolution detail present in the Fab, nanobody and bRIL fusion, but poorer resolution of 4-5 Å for SFXN1 in the detergent micelle. Local refinement, particle subtraction, and focused classification (fig. S2) improved the map to 3.4 Å, sufficient to build an atomic model for SFXN1 (fig. 1, also fig. S4).

**Figure 1:**
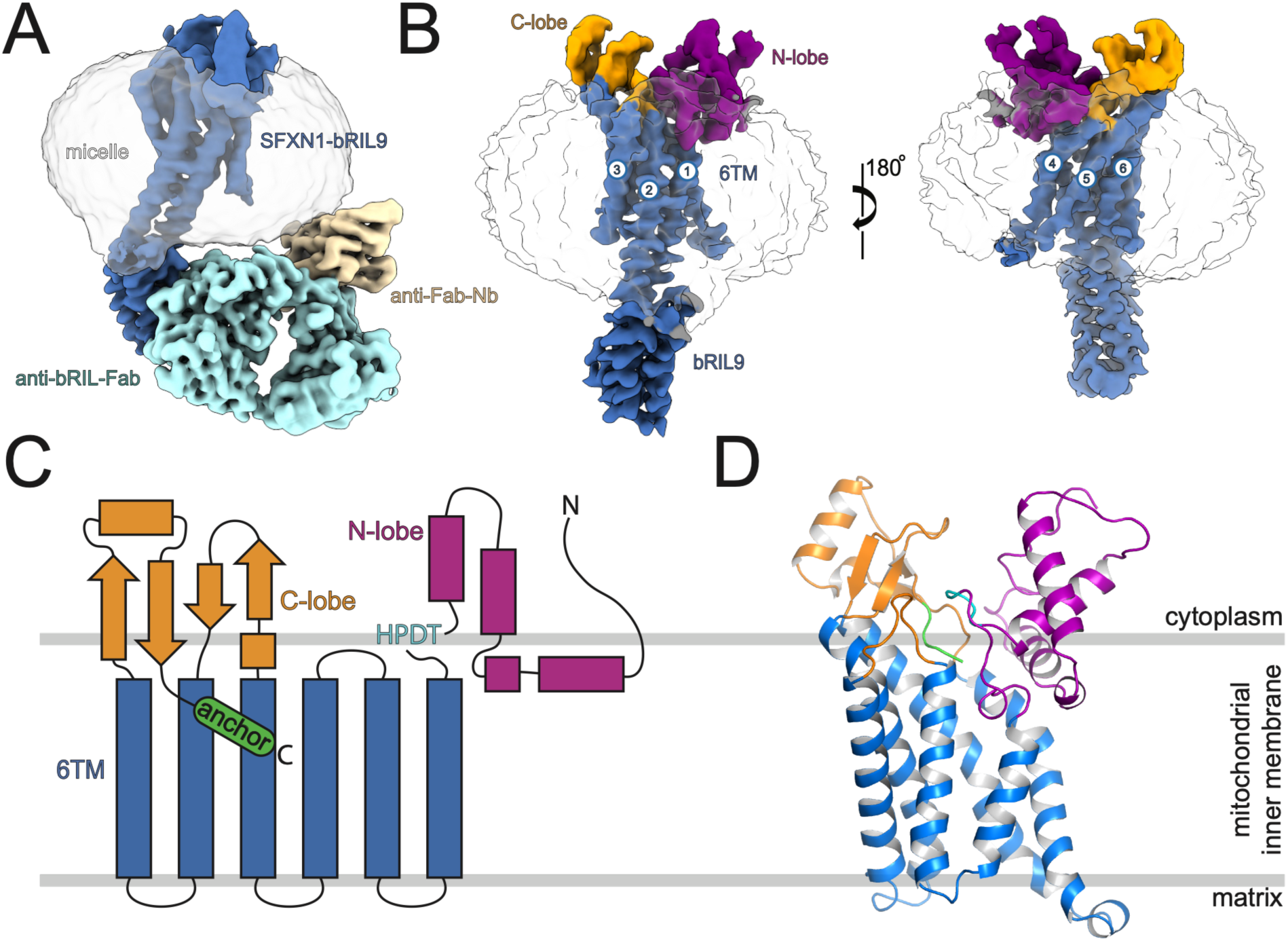
Cryo-EM structure of Human SFXN1. (**A**) Unsharpened cryo-EM map of the SFXN1-bRIL9:anti-bRIL Fab:anti-Fab nanobody complex in detergent. SFXN1-bRIL9 is coloured blue, anti-bRIL-Fab cyan, anti-Fab nanobody wheat, and the micelle transparent grey. (**B**) Sharpened (B-factor: - 90) cryo-EM map after focused refinement on SFXN1-bRIL9. Left, back view and right, front view, after a 180 ° rotation in the y axis. Circled numbers indicate TM helices. (**C**) Topology model of SFXN1 in the mitochondrial inner membrane. Rectangles indicate helices and arrows beta-sheets. The N and C-terminal lobes are coloured magenta and orange respectively. HPDT motif and the C-terminal anchor are cyan and green, respectively. (**D**) A cartoon rendering of SFXN1 coloured as in C, the bRIL is replaced with matrix loop 1 from an Alphafold2 model of SFXN1.

The structure reveals apo-SFXN1 as a monomer comprising six transmembrane (6TM) helices arranged in an asymmetric bundle and two soluble domains that form lobes on opposing sides of the bundle (fig. 1B, C and D). The N-terminal lobe (N-lobe; residues 1-93) contains four short helices (NH1-4) connected by short loops (fig. 1b-d). Preceding TM1 is a conserved HPDT motif (residues 78-89) (fig. 1B-D and fig. S5), which leads into a loop-helix-loop motif that runs parallel to the membrane outer leaflet. The C-lobe is derived from three discontinuous C-terminal segments (residues 135-144, 203-219, and 286-315; fig. 1C and D), with two pairs of anti-parallel beta-strands (CB1-4) buttressed by a solvent exposed helix (CH2) (fig. 1 and fig. S5). The C-lobe positions the C-terminus of the protein (residues 319-322) in the centre of the SFXN1 transmembrane helical bundle (fig. 1C and D). A structural similarity search identified no significant homology to proteins other than SFXN family members (*27*).

To determine the orientation of SFXN1 in the mitochondrial inner membrane, we employed an mNeonGreen (mNG)-based bimolecular fluorescence complementation system, in which fluorescence is reconstituted only when the two fragments are present in the same compartment (*28*). The large mNG(1–10) fragment was targeted to the mitochondrial matrix or intermembrane space by fusion to the N-terminal mitochondrial targeting sequences of COX8A or SMAC, respectively. Co-expression of SMAC-mNG(1–10) with mNG(11)-SFXN1 resulted in fluorescence reconstitution, whereas co-expression with COX8A-mNG(1–10) did not (fig. S6A-C). In contrast, mNG(11)-SHMT2, a control for mitochondrial matrix localization, reconstituted fluorescence with COX8A-mNG(1-10) but not with SMAC-mNG(1-10), confirming the specificity of the split-mNG system for sub-compartmental localization. Together with the experimental structure, these results establish that SFXN1 is positioned with its N- and C-lobes facing the intermembrane space, defining its topology in the mitochondrial inner membrane (fig. 1C and D).

### SFXN1 adopts a matrix-open conformation

SFXN1 has a solvent-accessible cavity that extends from the matrix side of the membrane to the approximate membrane midpoint (fig. 2A), whereas the cytoplasmic side lacks a comparable opening. On the cytoplasmic side, the N- and C-lobes are closed around the C-terminal anchor residues 319–322, specifically sandwiched by the conserved HPDT (residues 80–83) and FPQ (residues 287–289) motifs (fig. 2B–D and fig. S5).

**Figure 2:**
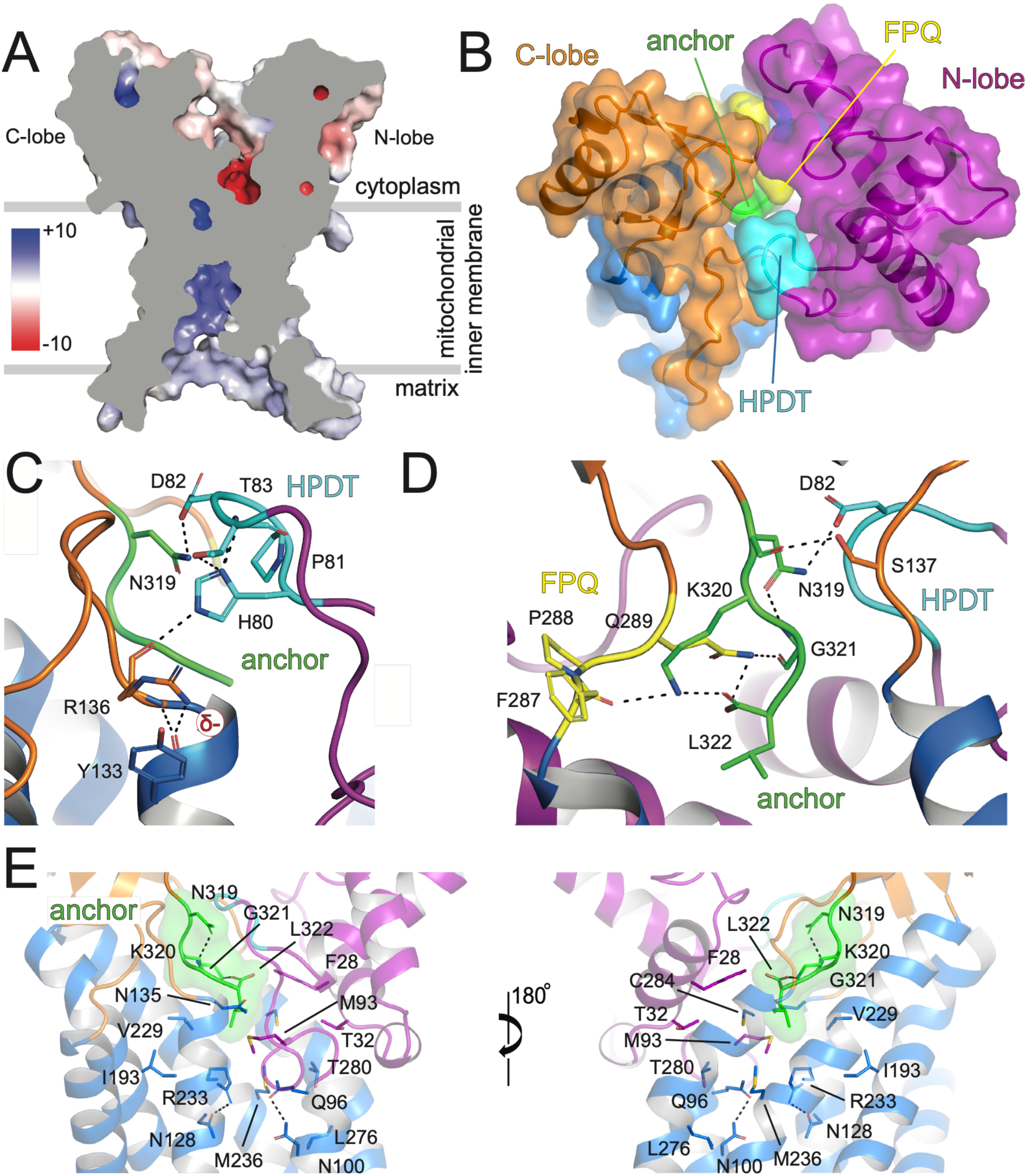
SFXN1 cryo-EM structure adopts a matrix open-cytoplasm closed state. (**A**) Surface rendering of SFXN1 showing a central slab through the protein. The electrostatic potential (kTe^-1^) was approximated using the APBS plugin in PyMOL and coloured from red (acidic) to blue (basic) from on a sliding scale from -10 to +10. (**B**) A cytosolic “top” view showing SFXN1 N and C-lobes with conserved elements buttressing the C-terminal anchor. (**C**) A close-up of the conserved HPDT motif in contact with the C-terminal end of TM2 and the C-terminal anchor residue N319. (**D**) A close-up of the conserved FPQ motif in contact with the C-terminal anchor. (**E**) Side views of the C-terminal anchor. The anchor is shown as a transparent green surface, above a cytosolic plug closing the central cavity to the cytosolic side of the membrane. The N-lobe is coloured purple, the C-lobe orange, the HPDT motif cyan, the FPQ motif yellow and the C-terminal anchor green. Side chains are coloured with CPK colouring. Dashed black lines indicate proposed bonding. The circled “δ-” symbol depicts a negative dipole moment at the C-terminal end of TM2.

The loop containing the HPDT motif adopts a horseshoe shape, with the HPDT residues forming the apex. The N-lobe HPDT motif contacts the C-lobe (fig. 2B), with a side chain interaction between D82 and N319 (fig. 2C-D). The HPDT motif also sits atop the C-terminal end of TM2, with H80 approaching within hydrogen bond distance of the open alpha helix at R136 backbone carbonyl, an interaction enhanced by the negative dipole at the C-terminal end of TM2. A cation-π interaction between R136 and Y133 also stabilizes the loop C-terminal to the TM2 helix (fig. 2C). On the other side of the C-terminal anchor, Q289 of the FPQ motif is hydrogen bonded to the G321 backbone carbonyl (fig. 2D), completing a FPQ motif: C-terminal anchor: HPDT motif sandwich. The side chain of the C-terminal residue, L322, rests amid a hydrophobic cluster that includes F28, M93, V229, M236, and C284 (fig. 2E), and the L322 C-terminal carboxylate forms a salt bridge back onto the C-terminal anchor residue K320 (fig. 2D). These residues sit atop two pairs of residues that make side chain contacts: N128 with R233 and Q96 with N100 (fig. 2E), collectively occluding the matrix-facing cavity of SFXN1 from the cytoplasm.

### Identification of SFXN1 leucine transporter activity

#### AlphaFold2 predicts a cytoplasmic-open state of SFXN1

A distinguishing feature of the SFXN1 matrix-open Cryo-EM structure is the closure of access to the cytoplasmic side of the membrane and the close approximation of the N- and C-lobes. It has been shown for GPCRs and transporters, that alternate states of receptors and transporters can be accessed by AlphaFold and similar models (*29*, *30*). AlphaFold2, run with default parameters, predicts a SFXN1 model with the same overall architecture of the matrix open state but instead features separated N- and C-lobes (fig. 3A). In the AlphaFold2 model, the matrix-facing cavity observed in the Cryo-EM structure is closed, whereas a cytoplasmic-facing opening extends toward the membrane midpoint (fig. 3B). Comparison of the matrix-open and cytoplasmic-open conformations shows that the D82 to N319 contact is no longer present (compare fig. 3C, left, to fig. 2C and D) in the cytoplasmic-open conformation. Furthermore, TM1 and TM2 are tucked into the matrix cavity, forming a matrix gate including hydrophobic residues—M107, M108, Y111, P181, M241, and V272—that are retracted in the matrix-open conformation but pack together in the cytoplasmic-open conformation, forming a hydrophobic barrier that occludes the substrate-binding site on the matrix side (fig. 3C, bottom). These differences suggest the presence of at least two discrete conformations, which may allow coordinated movements that couple closure of a matrix gate with opening of the cytoplasmic gate to allow access to a common substrate binding site.

**Figure 3:**
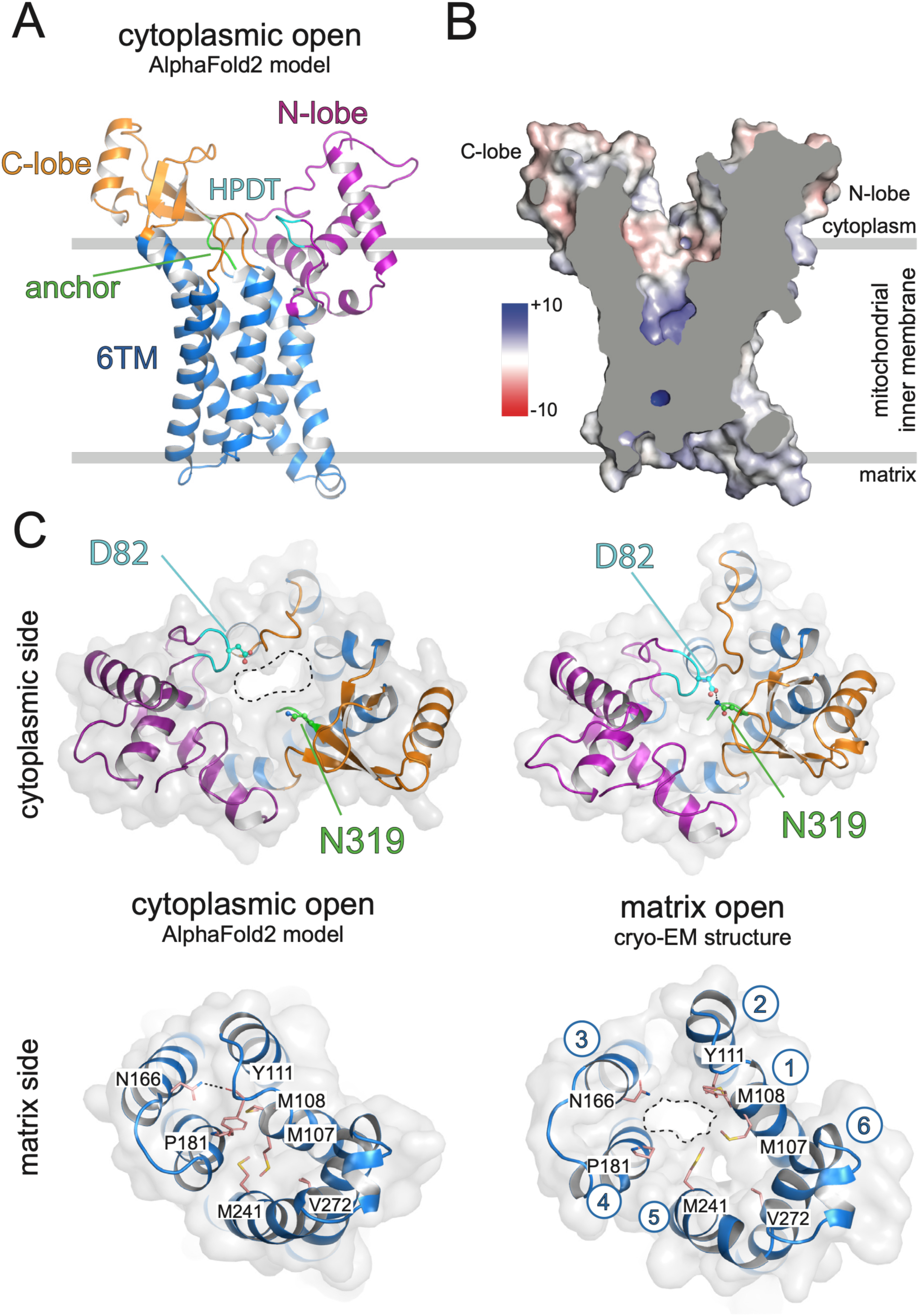
AlphaFold2 predicted SFXN1 structure adopts a cytoplasm open-matrix closed state. (**A**) Cartoon representation of SFXN1 as predicted by AlphaFold2 (SFXN1-AF2). The six-transmembrane helical bundle (6TM) is coloured blue and the N and C-terminal lobes are coloured magenta and orange respectively. The HPDT motif and the C-terminal anchor are cyan and green, respectively. (**B**) Surface representation of SFXN1-AF2 showing a central slab through the protein. The electrostatic potential (kTe-1) was approximated using APBS plugin in PyMOL and coloured from red (acidic) to blue (basic) on a sliding scale from -10 to +10. (**C**) Cytoplasmic open (AlphaFold2 model, left) and matrix open (cryo-EM structure, right) conformations viewed from the cytoplasmic (top) and matrix (bottom) side of the membrane. SFXN1 is rendered as a cartoon with a transparent grey surface and coloured as in fig. 1. A dashed black line encircles a central cavity exposed to the cytoplasmic side in the cytoplasmic open and in the matrix side in the matrix open conformations. Residues formative of the matrix gate are shown in pink. Transmembrane helices are numbered in the cryo-EM structure matrix side view. Selected residues are depicted as ball and sticks (top) and sticks (bottom) with CPK colouring.

#### Human metabolome-SFXN1 co-folding identifies hydrophobic amino acid binding

We leveraged AlphaFold3 and Boltz-2 (*31*, *32*) to predict potential SFXN1-ligand complexes using a human metabolite library (3,282 ligands from the Human Metabolome Database (*33*); fig. 4A). As an initial screen, we used AF3 to co-fold SFXN1 with the entire metabolite library using the matrix-open (fig. 1 and 2) and cytoplasmic-open (fig. 3) conformations as separate input models (custom templates), identifying 199 compounds that passed an initial confidence threshold (ligand pLDDT > 70 and ipTM > 0.8, fig. 4B) in both conformations (data S1). We next evaluated these candidates using Boltz-2 and assessed the binary affinity probability score as a confidence metric (fig. 4C). We observed a strong positive correlation (R^2^ = 0.72) between binary affinity probability scores in the matrix-open and cytoplasmic-open conformations, suggesting that the pocket structures in both forms are comparable despite their differences in cavity openness. The branched-chain amino acids leucine and isoleucine scored highly (fig. 4C), along with several other proteinogenic and non-proteinogenic amino acids (data S1).

**Figure 4:**
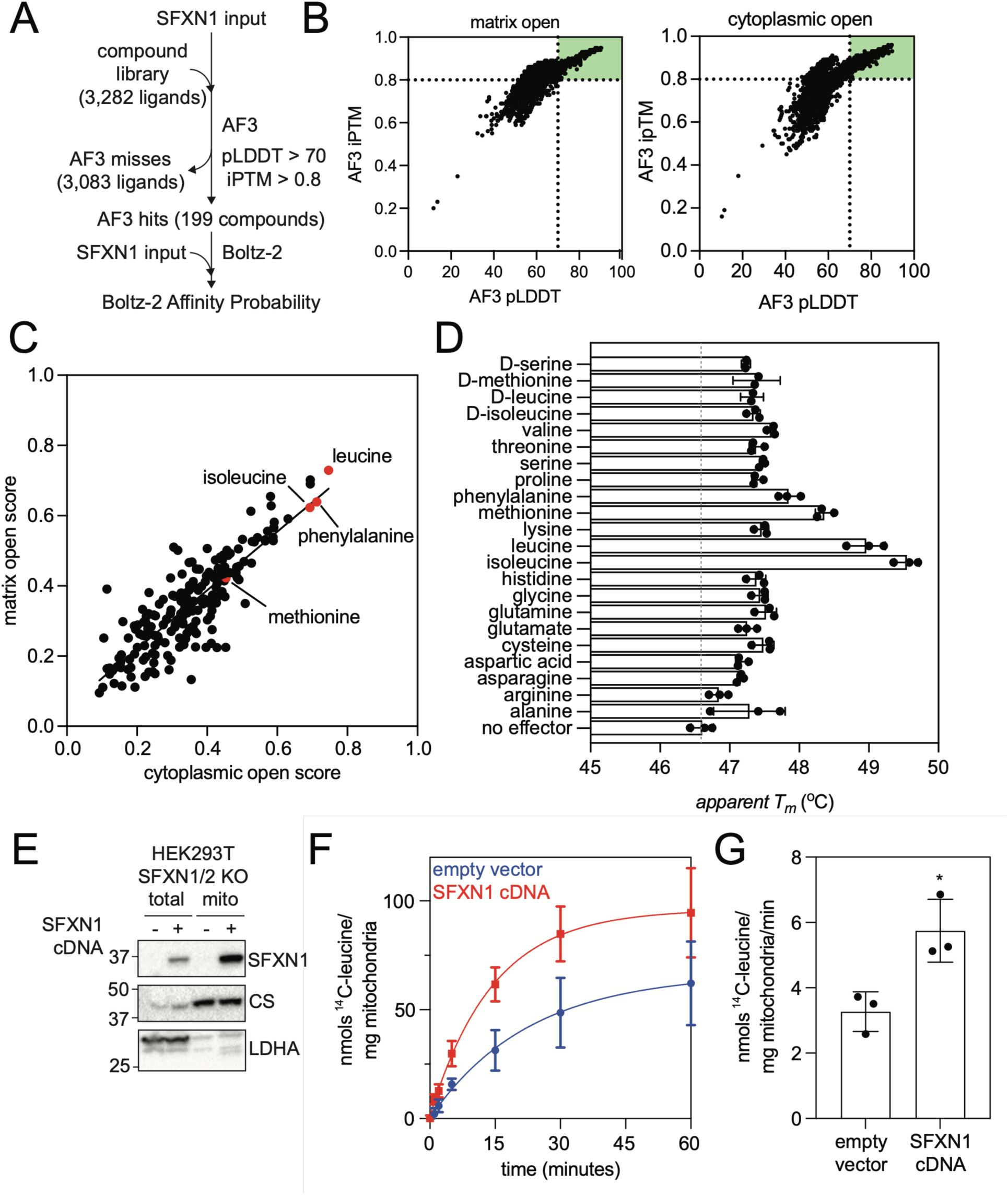
SFXN1 binds hydrophobic amino acids and transports leucine into the mitochondria. (**A**) Structural bioinformatics workflow for screening SFXN1 against a curated Human Metabolome Database compound library (see Methods). (**B**) AlphaFold3 outputs using the matrix open cryo-EM structure or the cytoplasmic open AlphaFold2 (AF2) model are used as inputs. The green box (pLDDT > 70 and iPTM > 0.8) indicates the compounds for subsequent Boltz-2 analysis. (**C**) Boltz-2 affinity probability scores using the matrix open cryo-EM structure or the cytoplasmic open AF2 model are used as inputs. Red circles indicate > 1 oC stabilisers in panel (d). (**D**) NanoDSF thermostability analysis of SFXN1 purified in detergent after incubation with 10 mM of indicated L-amino acid, or selected D- amino acids. The dashed grey line indicates the mean stability of the no effector control (**E**) Western blot of a representative mitochondrial isolation from HEK293T SFXN1/SFXN2 double knockout cells with SFXN1 cDNA addback or empty vector control. Citrate synthase (CS) and lactate dehydrogenase A (LDHA) serve as mitochondrial or cytoplasmic markers, respectively. (**F**) Time dependent 14C-leucine uptake in isolated mitochondria from (e). Data are presented as the mean of n=3 biological replicates with standard deviation shown as error bars. A one-phase association nonlinear model was fitted in Prism. (**G**) Initial uptake rates (0-5 mins) were determined for each biological replicate from (f) and plotted with the standard deviation shown as error bars. Student’s two-tailed T-test was used to compare the means between the initial rates of empty vector and SFXN1-cDNA addbacks (*P < 0.05).

Thermal stability analysis has been extensively utilised to probe transporter substrate interactions (*34–37*). We measured the thermal stability of SFXN1 in the presence of each proteinogenic L-amino acid (fig. 4D). Most amino acids gave small increases (<1 °C) in thermal stability on incubation at 10 mM. Stabilising effects larger than 1 °C were observed for leucine, isoleucine, methionine, and phenylalanine, which all ranked highly in Boltz-2 affinity probability scores in both states (fig. 4C). In contrast, the non-proteinogenic D-enantiomers of these molecules did not exhibit stabilization of SFXN1 beyond the general amino acid response, consistent with stereo selective interactions within the predicted pocket (fig. 4D).

#### SFXN1-dependent leucine uptake into isolated mitochondria

As leucine emerged as one of the strongest hits from both Boltz-2 analysis and thermal stability measurements, we next examined whether SFXN1 mediates mitochondrial leucine uptake. SFXN1 and SFXN2 are the predominant paralogues expressed in HEK293T cells and share high sequence similarity (*38*). To minimize potential functional redundancy, we generated a SFXN1/SFXN2 double knockout (DKO) line to minimize potential functional redundancy and stably reintroduced SFXN1 cDNA (*39*) (fig. S7B). We isolated mitochondria by differential centrifugation (fig. 4E) and measured radiolabelled ^14^C-leucine uptake (fig. 4F and G). Mitochondria isolated from SFXN1-rescued DKO cells exhibited a time-dependent increase in both total ^14^C-leucine uptake compared to DKO controls (fig. 4F) and an increased initial uptake rate measured over the first five minutes (fig. 4G). The increase in total uptake capacity indicates enhanced mitochondrial leucine accumulation and the increase in initial uptake rate is consistent with a direct, transport-mediated role for SFXN1 in mitochondrial leucine import.

#### Specific binding of branched-chain amino acids at a common substrate-binding site

Boltz-2 predicts that branched-chain amino acids occupy a common substrate-binding site, at the approximate membrane centre (fig. 5A). Independent of conformation, the hydrophobic side chain of both leucine and isoleucine pack against a hydrophobic surface formed by V185, A237, and L276 (fig. 5B). The amino and carboxylate group interactions differ subtly, but in each case these groups face a pocket of amide-rich residues that include Q96, N100, N124, N128 and N189, and R233 in the leucine bound matrix-open conformation. This binding site is supported by a high degree of conservation in TM1 (96-**<u>Q</u>**<τPM**<u>N</u>**M-101), TM2 (119-FWQI**<u>N</u>**Q[S/T]F**<u>N</u>**AVV-131) and TM4 (181-PFAA**<u>V</u>**AA**<u>N</u>**CIN-192) (binding site residues shown in bold and underlined). Because isoleucine exhibited the strongest stabilising effect in thermal shift assays, we used it as a representative amino acid to probe predicted binding interactions. We introduced point mutations at a set of predicted contact residues and assessed isoleucine-mediated stabilization of SFXN1 (fig. 5C). We found that N100A, N124A, N128A, N189A, and L276A abolished isoleucine-mediated stabilisation of SFXN1. In contrast, mutation of N192, a nearby but non-contacting residue, had no effect.

**Figure 5:**
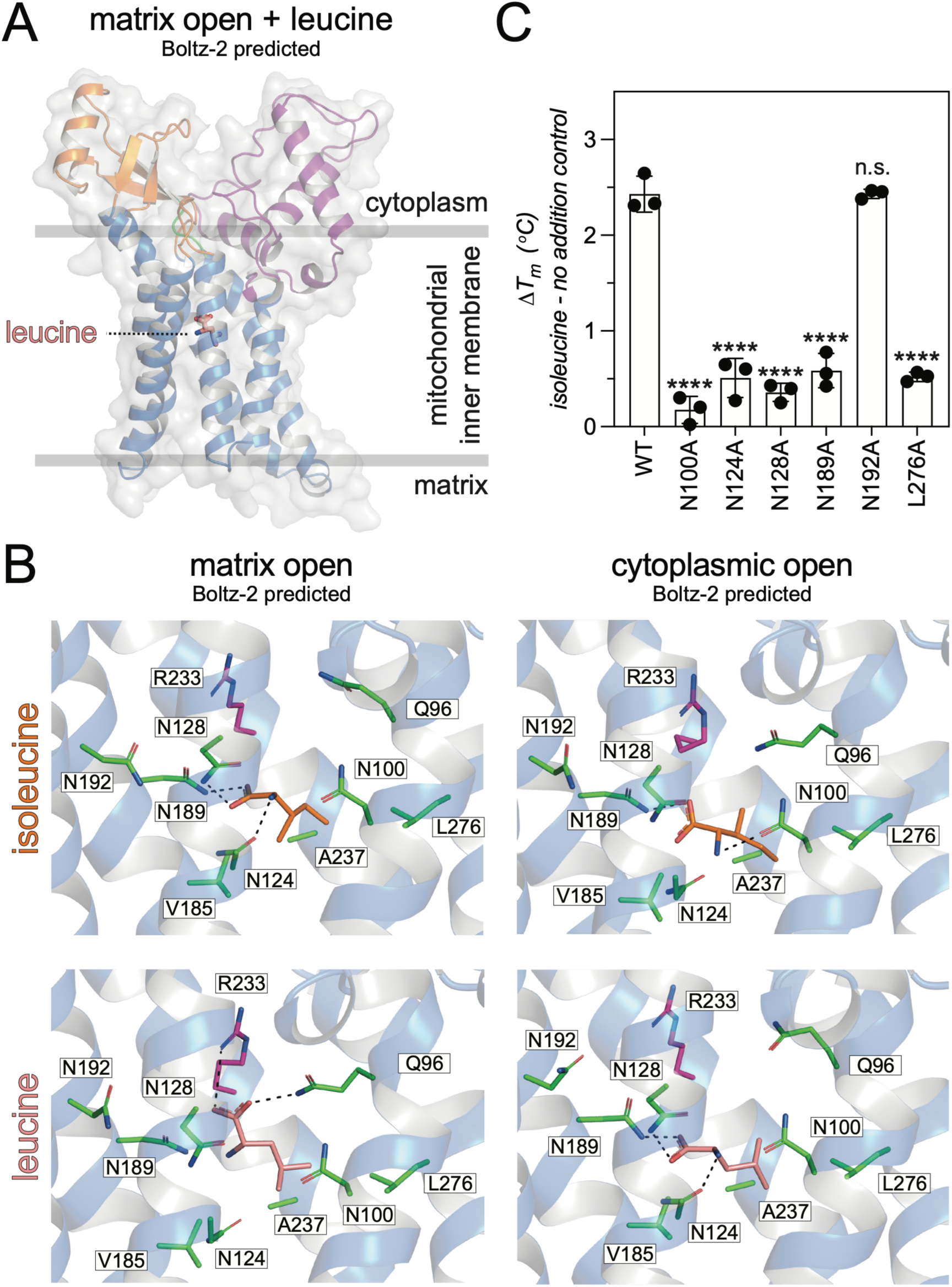
Site directed mutagenesis of the SFXN1 branched chain amino acid binding site. (**A**) Cytoplasmic view of the isoleucine (top) and leucine (bottom) bound substrate binding site after Boltz-2 prediction. AF2 cytoplasmic open conformation (left) and cryo-EM matrix open conformation (right) were used as input models. The cartoon backbone is coloured blue. Ligands and side chains are shown as ball-and-sticks. The isoleucine is shown as yellow, leucine cyan and residues in the substrate binding site are shown as green. Proposed hydrogen bonds are shown as black dashed lines. Atoms are shown with CPK colouring. (**B**) Side view of the matrix-open leucine bound conformation coloured as in (a) but with a transparent surface. (**C**) NanoDSF of purified SFXN1 wild type and substrate binding site mutant proteins. The ΔTm is calculated from the difference in protein thermostability upon the addition of 10 mM isoleucine or a no addition control. Data are presented as the mean of three biological replicates (n=3) with standard deviation shown as error bars. One-way analysis of variance (ANOVA) was used with Dunnett’s multiple-comparison post-hoc test to compare the difference between WT to all other samples individually (**P < 0.01;***P < 0.001;****P < 0.0001).

#### A C-terminal anchor toggle-switch mechanism

Sixty years ago, Jardetzky proposed that transporters operate through a common substrate-binding site flanked by alternately open gates on either side of the membrane (*40*). Comparison of the matrix-open and cytoplasmic-open conformations, and identification of a single substrate binding site at the approximate membrane center (fig. 5), suggests distinct cytoplasmic and matrix gates in SFXN1 that alternately control access to the substrate-binding site (fig. 3). To define the conformational changes underlying gate opening and closure across the SFXN1 protein, we calculated an alignment-free difference contact map between the matrix-open Cryo-EM structure and the cytoplasmic-open AlphaFold2 model (fig. S7). This analysis indicates that residues spanning the N-lobe through TM2 (residues 1-135) and TM4 through the C-terminal anchor (residues 171-322) move largely as opposing rigid bodies, with minimal intra-body rearrangement, but large rearrangements between bodies. For example, the distance changes between D82 and N319 (8 Å) and M108 and P181 (6 Å), respectively, report on interdomain movement between conformations (fig. 3C). TM3 and cytosolic loop 1 (residues 136-144), appear to not belong to either of these rigid bodies, and instead may serve as a linker connecting them (Movie 1, see fig. S7). Overall, the coordinated rigid-body movements of the N and C terminal bodies provide a structural basis for alternating exposure of the substrate-binding cavity, enabling reciprocal opening and closure of the matrix and cytoplasmic gates in a rocker-switch mechanism (fig. 2, 3 and 6).

**Figure 6:**
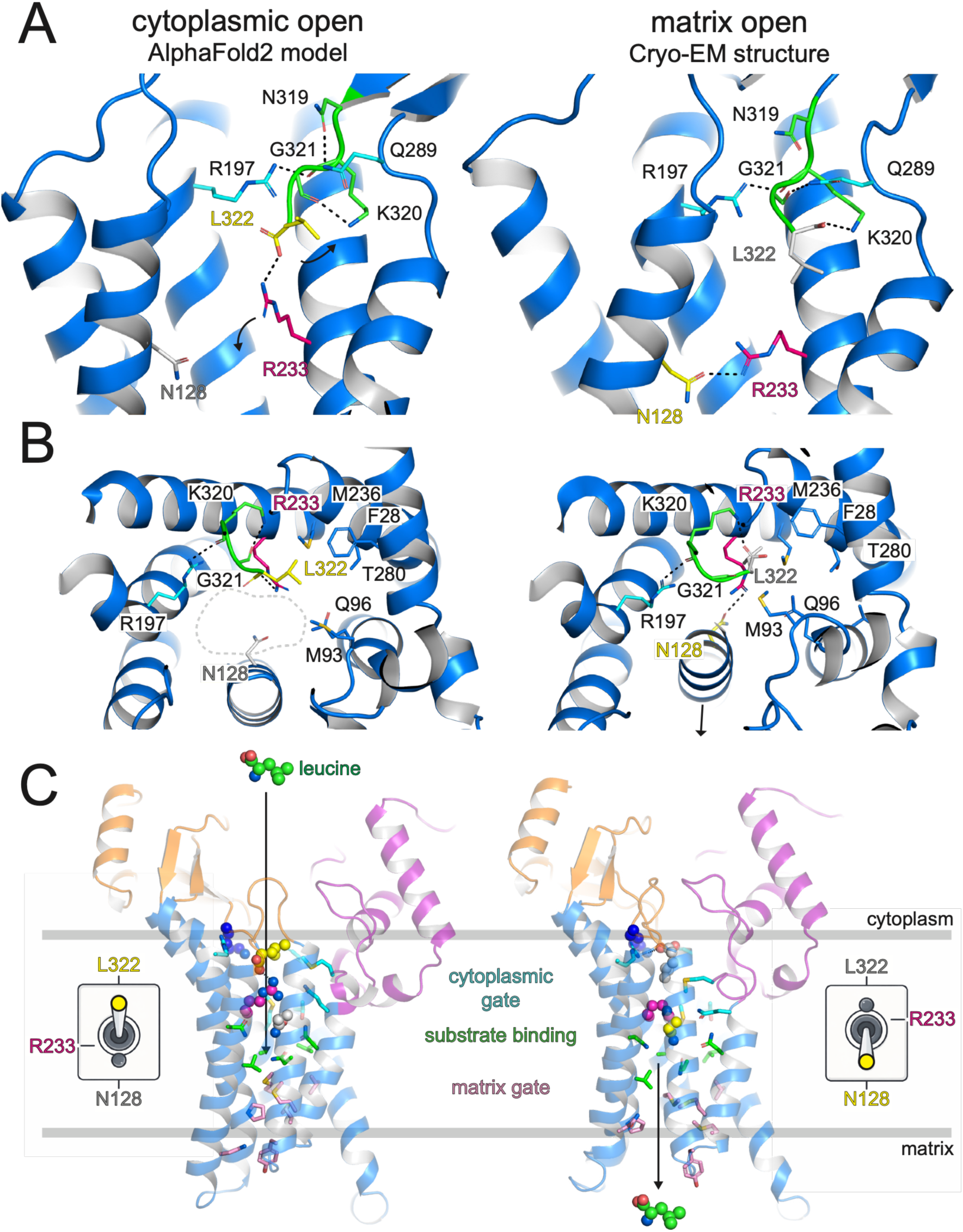
R233 toggle switch controls access to the substrate binding site. Cytoplasmic open (left) and matrix open (right) conformations of SFXN1, from AlphaFold2 prediction and Cryo-EM structure, respectively. (**A**) Side views of SFXN1 at the R233 toggle switch. The cartoon backbone is shown as blue, the C-terminal anchor is coloured green, R233 is coloured magenta, and neighbouring residues cyan. In the cytoplasmic open conformation N128 is coloured grey, and L322 is coloured yellow, and in the matrix open conformation N128 is coloured yellow, and L322 is coloured grey. Yellow indicating an interaction with R233, and grey not an interaction with R233 in that specific conformation. Curly arrows show predicted movements of R233 and L322 on transition from the cytoplasmic open to matrix open conformation. Dashed black lines indicated predicted hydrogen bonds. (**B**) Top view of (a), with additional residues plugging the cytoplasmic gate shown as blue sticks. A solid black arrow indicates outward movement of TM2 on transition from the matrix open to cytoplasmic open conformation. A dashed grey line indicates access to substrate binding site. (**C**) Side views of SFXN1 illustrating elements required for alternating access. SFXN1 backbone is coloured as in fig.1 with the N-lobe purple, the C-lobe, orange, and the 6TM blue. Cytoplasmic gate residues are coloured cyan (fig. 6b), substrate binding site residues green (fig. 5), and matrix gate residues pink and shown as sticks (fig. 3c). Residues N128, R233 and L322 are coloured as in fig. 6a, K320 is coloured blue. Leucine substrates is shown as green spheres, arrow indicates direction of transport. Residues are shown with with CPK colouring.

The cytoplasmic gate is centered on the C-terminal anchor residues K320, G321, and L322 (fig. 2, fig. 6A). In both conformations, the conserved R197 and Q289 sequester the C-terminal anchor by contacting K320 and G321 at the peptide backbone, respectively. In the cytoplasmic-open conformation, the L322 C-terminal carboxylate is predicted to interact with R233, and N128 is unbound, presumably awaiting substrate (fig. 5, fig. 6A and B, left). In contrast, N128 forms a hydrogen bond with R233 in the matrix-open conformation, whereas L322 forms a salt-bridge with K320 (fig. 6A and B, right, fig. S4H). R233 is therefore predicted to act as a toggle switch between conformations; it forms a salt-bridge with L322 in the cytoplasmic-open conformation, keeping access to the substrate binding site clear, and is predicted to engage N128 in the matrix-open conformation, capping the substrate binding site on the cytoplasmic side (Movie 2, see fig. 6).

Taken together, these observations indicate that access to the shared substrate-binding site is controlled by a cytoplasmic gate operating through a R233 toggle-switch mechanism, which allows crosstalk between the C-terminal anchor and the substrate binding site to control substrate access to the cytosolic side of the protein (fig. 6C).

## Discussion

Mitochondrial metabolism depends on the controlled exchange of metabolites and ions across an otherwise impermeable inner membrane. For decades, this exchange was largely attributed to members of the Solute Carrier Family 25 (SLC25), including the ADP/ATP carrier (*41*, *42*), the aspartate–glutamate carrier (*43–45*) and transporters for central organic acids (*46*). The subsequent identification of additional systems such as ATP-binding cassette transporters (*47–50*), the mitochondrial calcium uniporter (*51–53*), the mitochondrial pyruvate carrier (*54–59*) and the sodium–calcium exchanger NCLX (*60*, *61*) has revealed that inner-membrane transport is architecturally more diverse than originally appreciated.

The structure of SFXN1 presented here reveals a previously uncharacterized fold and captures a matrix-open conformation in which a central substrate-binding cavity is accessible from the mitochondrial matrix. Predictive modelling supports a complementary cytoplasmic-open state with access to the same substrate binding site, consistent with an alternating-access mechanism (*62*). Together, these structural and computational analyses support an alternating-access transport cycle in which leucine binds within a central cavity that is alternately exposed to either side of the membrane (*40*).

The substrate-binding cavity is not strongly charged, consistent with transport of an uncharged amino acid (fig. 2A and 3B). Instead, the cavity is enriched in polar amide residues positioned to coordinate the amino and carboxyl groups of the substrate, while an opposing hydrophobic surface accommodates hydrophobic side chains (fig. 5 and 6). Conserved motifs contribute to this hydrogen-bonding network, and their preservation from mammals to yeast and plants suggests that recognition and perhaps transport of hydrophobic amino acids into mitochondria by SFXN1 is evolutionarily ancient through Eukarya (fig. S5).

Previous work proposed that SFXN1 functions as a mitochondrial serine transporter(*20*). The studies reported here, however, revealed hydrophobic amino acids as providing the greatest stabilization of SFXN1 in thermal shift assays. Moreover, in our structural bioinformatics analysis leucine and isoleucine scored most highly for their compatibility with both matrix-open and cytoplasmic-open states, and mitochondrial uptake assays demonstrate robust SFXN1-dependent mitochondrial leucine uptake. Together, these observations confirm that SFXN1 can carry out mitochondrial leucine import, consistent with the structural features of the binding cavity. Nevertheless, it remains possible that SFXN1 can interact with and transport other amino acids in a context-dependent manner. Substrate selectivity may depend on association with modulatory proteins or other factors, metabolic context, transport mode (uniport versus exchange), counter-substrate availability, membrane potential, or assay-dependent parameters. Resolution of the full substrate spectrum will require systematic transport measurements in a fully reconstituted system under defined driving forces.

SFXN1 is monomeric (fig. 1 and fig. S1) and accomplishes its transport cycle with a small mass (∼32 kDa), comparable in scale to canonical mitochondrial carriers, yet operates without internal symmetry (*63*). Given the robust SFXN1-dependent leucine uptake observed in isolated mitochondria (fig. 4), we propose a rocker-switch mechanism in which coordinated rigid-body movements at the N and C-terminal ends of the protein alternately open cytoplasmic and matrix gates through a R233 toggle-switch mechanism, allowing leucine release into the matrix. Changes in conformations may be driven by binding of substrate, such as the predicted interaction of leucine with R233 and Q96 in the matrix-open conformation (fig. 5B), but this will require further investigation.

Public expression datasets and prior literature indicate that sideroflexins are broadly expressed across human tissues, including liver, kidney, and brain (*64–66*), with SFXN1 enriched in the periportal zone of the mouse liver—the primary site of hepatic amino acid catabolism (*67*, *68*). While SLC25A44 has been implicated in leucine metabolism in brown adipose tissue, its role in other tissues remains unclear (*69*). Whether SFXN1 and SLC25A44 perform overlapping or distinct roles in mitochondrial leucine transport remains to be determined.

More broadly, this study illustrates how integration of Cryo-EM structure determination, structural bioinformatics, and targeted functional assays can uncover the substrates and mechanisms of previously uncharacterized membrane transporters. Given the links between branched-chain amino acid metabolism and metabolic disease, defining the molecular basis of mitochondrial leucine transport establishes a foundation for understanding how amino acid flux across the mitochondrial inner membrane shapes cellular metabolism and physiology.

## Acknowledgements

We thank Drs. Rachelle Gaudet, Andrew Kruse, and all members of the Kory lab for helpful discussion. The Centre of Macromolecular Interactions at Harvard Medical School. Prof. Alan Brown for review of the manuscript and assistance with map interpretation. Dr. Shaun Rawson for advice in Cryo-EM data processing. Dr. Richard Walsh and the Molecular Electron Microscopy Suite at Harvard Medical School for Cryo-EM data collection. Dr. Meredith Skiba and the Kruse lab for bRIL fiducial markers.

## Data, Code, and Materials Availability

All data and code needed to evaluate and reproduce the results in the paper are present in the paper and/or the Supplementary Materials.

## Author Contributions

F.M.F., D.T.D.J., S.C.B., and N.K. designed the study and experiments. N.K., E.R.S.K. and S.C.B. acquired funding. D.T.D.J. processed cryo-EM data and built atomic models with assistance from E.R.S.K. D.T.D.J. and F.M.F. wrote the manuscript. F.M.F., D.T.D.J., S.C.B., and N.K. edited the manuscript. N.P. performed protein design of SFXN1-bRIL. B.F. conducted AF3 and Boltz-2 analyses. F.M.F., D.T.D.J, H.A., E.K., S.J., L.E.V. and W.H.T. performed experiments. S.A.A. and S.B. prepared reagents. S.A.A. established and implemented the split fluorescent complementation assay. All authors reviewed and approved the manuscript.

