## Supplementary materials for "Molecular basis of mitochondrial leucine transport by human Sideroflexin 1"

### Materials and methods:

#### Split-mNeonGreen bimolecular fluorescence complementation assay

Coding sequences for SMAC–mNeonGreen(1–10) and COX8A–mNeonGreen(1–10), comprising the first 40 or 25 amino acids of human SMAC/DIABLO or COX8A, respectively, as well as codon-optimized mNeonGreen(11)–SFXN1 and SHMT2–mNeonGreen(11), were cloned into pMXs-IRES-blasticidin (Addgene #83356).

Lentivirus was produced in HEK293T cells by co-transfection of transfer plasmids with pCMV-Gag-Pol (Addgene #8449) and pCMV-VSVG (Addgene #8454) using X-tremeGENE 9, according to the manufacturer's instructions. Viral supernatant was collected, filtered (0.45 µm), and used for transduction. Stable cell lines co-expressing split mNeonGreen components were generated by sequential transduction with mitochondrial-targeted mNeonGreen(1–10) constructs followed by mNeonGreen(11)-tagged proteins.

For imaging, cells were seeded onto poly-D-lysine–coated coverslips, fixed with 3% paraformaldehyde and 0.1% glutaraldehyde, permeabilized in 2% BSA and 0.1% Triton X-100, and stained with Alexa Fluor 594–conjugated anti-TOMM20 antibody (Abcam, 1:300). Coverslips were mounted using ProLong Gold antifade reagent (Invitrogen). Images were acquired using an Abberior Facility Line STED microscope with a 100× oil immersion objective. mNeonGreen fluorescence was quantified in 40 randomly selected cells per condition using ImageJ, applying a uniform threshold and measuring integrated intensity per cell. For flow cytometry analysis, cells were trypsinized and resuspended in PBS. A total of 50,000 events per sample were acquired on a BD LSRFortessa cell analyzer (BD Biosciences) with mNeonGreen excited at 488 nm. Data were analyzed using FlowJo (Becton Dickinson), and the mean mNeonGreen fluorescence intensity for each group was tabulated.

**Construction of *S. cerevisiae* expression strains**

Human SFXN1 cDNA was codon-optimized for expression in *S. cerevisiae*. A 3x FLAG tag followed by a 3C protease cleavage site was introduced at the N-terminus. The coding sequence was cloned into a modified *S. cerevisiae* expression vector, pDDGFP2-LEU2d, under control of an inducible galactose promoter[70]. Expression vectors were transformed into the *S. cerevisiae* BJ5460 strain (MATa ura3-52 trp1 lys2-801 leu2Δ1 his3Δ200 pep4::HIS3 prb1Δ1.6R can1 GAL) using the LiAc/SS carrier DNA/PEG method[71]. Transformed cells were selected on synthetic complete medium lacking uracil (SC-Ura; Merck) supplemented with 2% glucose. DNA fragments were amplified by polymerase chain reaction using Phusion polymerase (NEB) following the manufacturer's instructions.

***S. cerevisiae* expression of SFXN1-bRIL9 and purification**

Six hundred mL of preculture in SC-Ura medium supplemented with 2% glucose media was used to inoculate 12 L of YPG + 0.1% glucose to an $OD_{600}$ of 0.3. Cells were grown at 27 °C until an $OD_{600}$ of 0.6 was reached and induced with 2% galactose for 20 hours. The culture was harvested the following day by centrifugation (3000 x g, 20 min, 4 °C), washed in ice-cold Milli-Q water, and centrifuged under the same conditions. The cells were resuspended in 250 mL breaking buffer (100 mM Tris-HCl pH 8.0, 5 mM EDTA, 5 mM amino hexanoic acid, 5 mM benzamidine) supplemented with cOmplete Mini, EDTA-free Protease Inhibitor Cocktail tablets (Roche) and lysed mechanically by passing the suspension five times through a high-pressure homogenizer (AH-NANO, Duoning Biotechnology Group) at 1300-1450 bar pressure. Whole cells and debris were removed by centrifugation (3000 x g, 20 min, 4 °C) and the membranes were collected by ultracentrifugation (205,100 x g, 1 hr, 4 °C). Pellets were resuspended in wash buffer (100 mM Tris-HCl (pH 7.4), 5 mM amino hexanoic acid, 5 mM benzamidine) and collected as before. The final pellet was resuspended to approximately 20 mg $ml^{-1}$ in HBG buffer (20 mM HEPES (pH 7.4),

150 mM NaCl, 10% glycerol), snap-frozen in liquid nitrogen and stored at −80 °C until use. Protein concentration was determined using a Bicinchoninic acid (BCA) protein assay kit (Thermo).

Isolated membranes (1 g) were solubilized in 1.5% (w/v) lauryl maltose neopentyl glycol (Anatrace) with 20 mM HEPES (pH7.4), 150 mM NaCl and two cOmplete Mini EDTA-free protease inhibitor tablets (Roche) for 1 hour at 4 °C. The lysate was clarified by centrifugation (205,100 x g, 1 hour, 4 °C) before applying to Anti-FLAG Affinity Isolated Mouse Monoclonal Antibody M2 resin (Merck). The resin was washed with 25 column volumes of wash buffer (20 mM HEPES (pH 7.4), 150 mM NaCl, 0.1 mg $ml^{-1}$ tetraoleoyl cardiolipin (Avanti Polar Lipids) and 0.1% lauryl maltose neopentyl glycol) under gravity flow. The protein was eluted using 3x FLAG tag peptide (Merck) in elution buffer (20 mM HEPES (pH 7.4), 150 mM NaCl, 0.1 mg $ml^{-1}$ tetraoleoyl cardiolipin). The FLAG peptide was removed with a PD10 desalting column (GE Healthcare) and subsequently concentrated using a 100 kDa MWCO centrifugal concentrator (Amicon), snap-frozen in liquid nitrogen and stored at –80 °C.

**Size exclusion chromatograph with multi-angle laser light scattering (SEC-MALS)**

SFXN1 was injected onto a size exclusion column (Superdex 200 10/300 GL Increase) prior to SEC-MALS. Experiments were performed with a Wyatt Dawn Heleos II multi-angle light scattering detector with an Optilab T-rEX refractive index monitor and Agilent isocratic HPLC system equipped with a SEPAX SRT C SEC-300 column. 100 µL of 30 µM BSA or SFXN1 (50 mM HEPES (pH 7.4), 150 mM NaCl, 0.005 mg $ml^{-1}$ tetraoleoyl cardiolipin and 0.005% lauryl maltose neopentyl glycol), was injected at a flow rate of 0.4 mL/min. Peak alignment, band broadening, light scattering detector normalization were performed on the monodispersed BSA monomer peak. Data analysis was performed with the Astra software package version (version 7.3.2.21).

**SFXN1-bRIL9 for single particle Cryo-EM**

For structural studies, the protein was thawed and 20 µg HRV 3C protease from HRV 3C protease solution kit (Pierce) was added to remove the 3x FLAG tag at 4 °C overnight with rotation. The protein mixture was centrifuged at 21,000 g for 15 min at 4 °C, and subsequently passed through a Durapore® PVDF 0.1 µm filter column (Millipore). The protein sample was injected onto a Superdex S200 10/300 GL column equilibrated in SEC buffer (20 mM HEPES (pH 7.4), 150 mM NaCl, 0.005 mg $ml^{-1}$ tetraoleoyl cardiolipin (Avanti Polar Lipids) and 0.005% lauryl maltose neopentyl glycol). The peak fraction was collected and concentrated using a 100 kDa MWCO centrifugal concentrator to ~2.5 mg/mL.

SFXN1-bRIL9 was mixed with 1.1-fold molar excess of Anti-bRIL Fab [28–30], and 2-fold molar excess of anti-Fab nanobody and incubated on ice overnight. The complex was further purified by size exclusion chromatography (Superdex S200 Increase 10/300) equilibrated in SEC buffer (20 mM HEPES (pH 7.4), 100 mM NaCl, 0.005 mg $ml^{-1}$ tetraoleoyl cardiolipin (Avanti Polar Lipids) and 0.005% lauryl maltose neopentyl glycol). Peak fractions were concentrated using a 100 kDa MWCO centrifugal concentrator to ~1.8 mg/mL

**Cryo-EM data acquisition and processing**

The complex was vitrified on UltrAuFoil® holey carbon grids (Gold R 1.2/1.3 300 mesh size) using a FEI Vitrobot Mark IV (FEI, Hillsboro). Grids were glow discharged, and 3 µL of sample loaded to the grid in a chamber at 22 °C and 100% humidity. Samples were applied at a force of 15 and blotted for 6-8 seconds before plunge freezing in liquid ethane.

Data were collected using a ThermoFisher Scientific Titan Krios operated at 300 kV with a Slectris Energy Filter and Falcon4i direct electron detector in counting mode with a total exposure dose of ~51 $e^-/Å^2$. ThermoScientific Smart EPU software was used for acquisition. 82 frames per movie were collected at a nominal magnification of 165,000x, corresponding to 0.74 Å/pixel. Micrographs were collected at a defocus of -0.8 to -2.0 µm. The data processing scheme is summarised in supplementary Fig. 2; In brief, a CryoSPARC[72] Live session was established with patch motion correction followed by patch CTF estimation. We subsequently proceeded with a full data collection comprising 16,344 micrographs given automated particle picking, followed by 2D classification, *ab initio* reconstruction, and refinement, produced a map displaying features of the SFXN1–bRIL9 complex.

Initial particle picking using the blob picker (150 Å diameter) yielded 2,116,705 particles, which were extracted with a 256 Å box size and Fourier cropped to 128 Å. We performed 2D classification and selected a broad set of particle views exhibiting clear SFXN1 complex features to generate an *ab initio* model. Particles that were not selected were used to generate three junk classes. We then performed four sequential rounds of heterogeneous refinement and *ab initio* reconstruction, resulting in a particle stack of 282,282 particles (supplementary Fig. 2, "A–D"). Non-uniform refinement yielded a map at 3.77 Å resolution, which was used to generate templates for template-based particle picking.

A CTF resolution cutoff of <6 Å was applied, rejecting 2,805 exposures, and template picking subsequently extracted 7,459,368 particles using a 360 Å box size Fourier cropped to 160 Å. Sequential rounds of *ab initio* reconstruction and heterogeneous refinement ("E-H") reduced this dataset to a particle stack of 428,875 particles, which upon non-uniform refinement yielded a map at 3.37 Å resolution.

Particles were then re-extracted with a 360 Å box size and 256 Å Fourier crop, and non-uniform refinement was repeated with estimation of per-particle-scale, improving the resolution to 3.25 Å. We next performed rebalance orientations followed by subset particles based on their per-particle scale values, reducing the particle stack to 229,197 particles with reasonably balanced orientations and good particle quality as judged by the per-particle scale metric. We then performed reference-based motion correction resulting in an unbinned particle stack with 360 Å box size.

Subsequent non-uniform refinement followed by two rounds of global CTF refinement produced a map at 3.15 Å resolution after homogeneous reconstruction, hereafter referred to as the "global map." This map displayed excellent features for the anti-bRIL Fab and anti-Fab nanobody (fiducial markers); however, the map for SFXN1 within the detergent micelle remained comparatively poor (supplementary Fig. 3).

We therefore adopted a strategy to focus refinement on SFXN1 while reducing the contribution from bRIL and the fiducial markers. To perform particle subtraction of the fiducial markers, we generated a mask surrounding these regions and performed local refinement. The same mask was then used to subtract particles within this masked region. Reconstruction of the resulting particle stack still showed residual density corresponding to fiducial markers; therefore, particles were realigned and the subtraction procedure was repeated once.

We used the fiducial marker subtracted particle stack and a mask around SFXN1 protein alone as inputs to an initial local refinement. This map had enhanced resolution for SFXN1, however, artifacts such as radial spokes originating from the detergent micelle were observed. To address this, we generated a mask encompassing SFXN1–bRIL together with the detergent micelle (personal communication from Dr. Shaun Rawson, HMS) and performed local refinement. This produced a map at 3.34 Å resolution with improved density for SFXN1, without such artifacts.

Despite these improvements, the map still displayed poorer resolution in the region corresponding to the C-lobe. We therefore generated a mask surrounding this region and performed 3D classification without alignment, which yielded improved density for this portion of the protein ("H"). The final reconstruction was obtained from 147,376 particles, yielding a map at a reported resolution of 3.34 Å. The final map is shown after micelle estimation and subtraction with LocScaleSurfer plugin in ChimeraX[73,74].

**Model building and refinement**

We used the AlphaFold2[7]-predicted structure of SFXN1-bRIL9 as an initial model for model building. The model was manually fitted into the map in ChimeraX, and ISOLDE[73–75] with local distance constraints to improve agreement with the map. Iterative cycles of manual model building in Coot and ISOLDE[75,76] and refinement using Servalcat (REFMAC) [76] were performed. The final model was evaluated by MolProbity[77]. Statistics of map reconstruction and model refinement are presented in supplemental table 1. Figures were prepared using PyMOL (Version 3.0, Schrodinger LLC) and ChimeraX[73,74].

**Human cell culture for site directed mutagenesis studies**

For substrate binding site investigations, SFXN1 wildtype and variants were expressed in HEK293T cells. HEK293T SFXN1 cells were cultured at 37 °C in a humidified atmosphere containing 8% (v/v) $CO_2$. Cells were passaged in Dulbecco's Modified Eagle's Medium (DMEM) supplemented with 10% (v/v) fetal bovine serum (FBS), 1% (v/v) penicillin–streptomycin (Pen/Strep), and 1% (v/v) glutamine until confluent. Confluent cultures were detached using trypsin and transferred to 3 L flasks containing 1 L Freestyle medium supplemented with 1% (v/v) Pen/Strep and 1% (v/v) FBS. Cultures were incubated until cells reached a density of approximately 1–3 × $10^6$ cells $mL^{-1}$. Cells were harvested (3000 x g, 20 min, 4 °C), and the cell pellet was washed in phosphate-buffered saline (PBS) and stored at -80 °C.

**Thermal stability measurements using differential scanning fluorimetry**

Thermal unfolding analysis was performed using dye-free differential scanning fluorimetry[78]. Approximately 5 µg of protein was added into a final volume of 12 µL of buffer (20 mM HEPES (pH 7.4), 150 mM NaCl, 0.1 mg $ml^{-1}$ tetraoleoyl cardiolipin) and, when required, 1 or 10 mM compound. The samples were loaded into nanoDSF-grade standard glass capillaries. The temperature was increased by 5 °C every minute from 20° to 95°C, the intrinsic fluorescence was measured in a Prometheus NT.48 nanoDSF device, and the apparent $T_m$ was calculated with the PR.ThermControl software (NanoTemper Technologies). A $\Delta T_m$ value was obtained by subtracting the $T_m$ of the protein in absence of effectors from the $T_m$ on the addition of effectors.

**Metabolite library preparation for co-folding**

To identify potential endogenous ligands of SFXN1, we developed a structure-based virtual screening workflow leveraging complementary protein–ligand co-folding tools (AlphaFold3 and Boltz-2). We assembled a library of 3,282 candidate ligands with experimental quantification and at least 5 heavy (non-hydrogen) atoms from the Human Metabolome Database (HMDB). To adjust the SMILES strings from the neutral forms retrieved from the HMDB, we used OpenBabel (version 3.1.0) to predict the dominant protonated form of all ligands at pH 7.0 and used the OpenBabel SMILES string outputs in all subsequent co-folding. Two template structures representing the cytoplasmic-open (AF2 model) and mitochondrial matrix-open (cryo-EM structure) conformational states of the transporter were used with the same virtual screening methodology as described below.

**Virtual screening methodology for protein-ligand co-folding**

We first performed protein–ligand co-folding using AlphaFold3 (version 3.0.1). We ran a single initial prediction of the SFXN1 amino acid sequence in the absence of any ligand to generate a multiple sequence alignment (MSA) using the default inference pipeline. We used the same resulting MSA and provided exactly one template structure at a time as input to all AlphaFold3 (AF3) co-folding. AF3-predicted structures were generated using 10 recycles and 5 diffusion samples for each of the 3,282 candidate ligands for both the cytoplasmic-open and matrix-open template structures. The top-ranked diffusion sample for each AF3 output as determined by the AF3 “ranking_score” was evaluated for ligand pLDDT > 0.7 and ipTM > 0.8. Compounds that passed both thresholds were carried forward into subsequent Boltz-2 analysis. This filtering identified 325 hits for the cytoplasmic-open template and 343 for the matrix-open template.

To confirm and rank the hits identified by AF3, we used the “affinity_probability_binary” metric computed by Boltz-2 (version 2.2.1). As with the AF3 input preparation, we generated a single MSA for the SFXN1 sequence using MMseqs2 through the ColabFold MSA server following the default Boltz-2 input preparation pipeline. Boltz-2 predicted structures were generated using the default Boltz-2 inference parameters (3 recycles, 1 diffusion sample, 200 diffusion steps) for each of the candidate ligands passing the AF3-screen confidence thresholds. We again evaluated the cytoplasmic-open and matrix-open structures by providing them as templates for the co-structure prediction one at a time with template forcing enabled (using an RMSD threshold 1.5 Å) along with inference potentials (--use_potentials flag) and molecular weight correction for the affinity prediction module (--affinity_mw_correction flag).

**Mitochondria isolation**

HEK293T cells were cultured in FreeStyle® media between 0.5 and 3 million cells/mL in 1 L baffled flasks with continuous shaking (37 °C, 8% $CO_2$, 80 rpm). The evening before isolation, 150 mL of cells were split to 1 million cells/mL into a 0.5 L flask and incubated overnight with shaking. Cells were harvested the next morning and washed twice in ice cold PBS without calcium and magnesium chloride. All following steps were carried out on ice and using wide-bore P1000 pipette tips. Cell pellets were resuspended in mitochondria isolation buffer (MIB, 300 mM sucrose, 10 mM HEPES, 0.2 mM EDTA, 0.1% fatty-acid free BSA, pH 7.4) and dounce homogenised with 20 passages and centrifuged at 700 x g for 5 minutes. The resulting supernatant was removed and stored, whilst 1 mL of MIB was used to resuspend the pellet. The resuspended pellet was dounce homogenised with 20 passages and centrifuged at 700 x g for 5 minutes. The supernatant was removed and combined with the previously collected supernatant and centrifuged at 10,000 x g for 10 minutes. The resulting mitochondrial pellet was resuspended in MIB without bovine serum albumin and centrifuged at 10,000 x g for 10 minutes. The resulting pellet was resuspended in mitochondrial assay buffer (MAB, 70 mM sucrose, 220 mM mannitol, 10 mM $KH_2PO_4$, 5 mM $MgCl_2$, 2 mM HEPES, 1 mM EGTA, 0.2% fatty acid free BSA, pH 7.4 (KOH)), and BCA assay performed to determine protein content. Protein was adjusted to 1.5 mg/mL before uptake assays were performed.

**Mitochondria uptake assays**

20 μL of mitochondria in MAB buffer were pipetted to the bottom of a 1.5 mL Eppendorf tube. The reaction was initialised by the addition of 80 μL of MAB buffer + 6 μM of $^{14}C$-Leucine at room temperature (Cambridge Isotope Laboratories inc., MA). Uptake was quenched by the addition of 1 mL of ice-cold MAS buffer, followed by rapid filtration and washing with three 1 mL additions of MAB buffer on mixed cellulose ester membranes (0.45 μm) using a Hoeffer vacuum manifold. Membranes were collected and moved to scintillation vials, before addition of 4 mL of scintillant (EcoLume, MP Biomedicals) and radioactivity quantified on a Hidex scintillation counter.

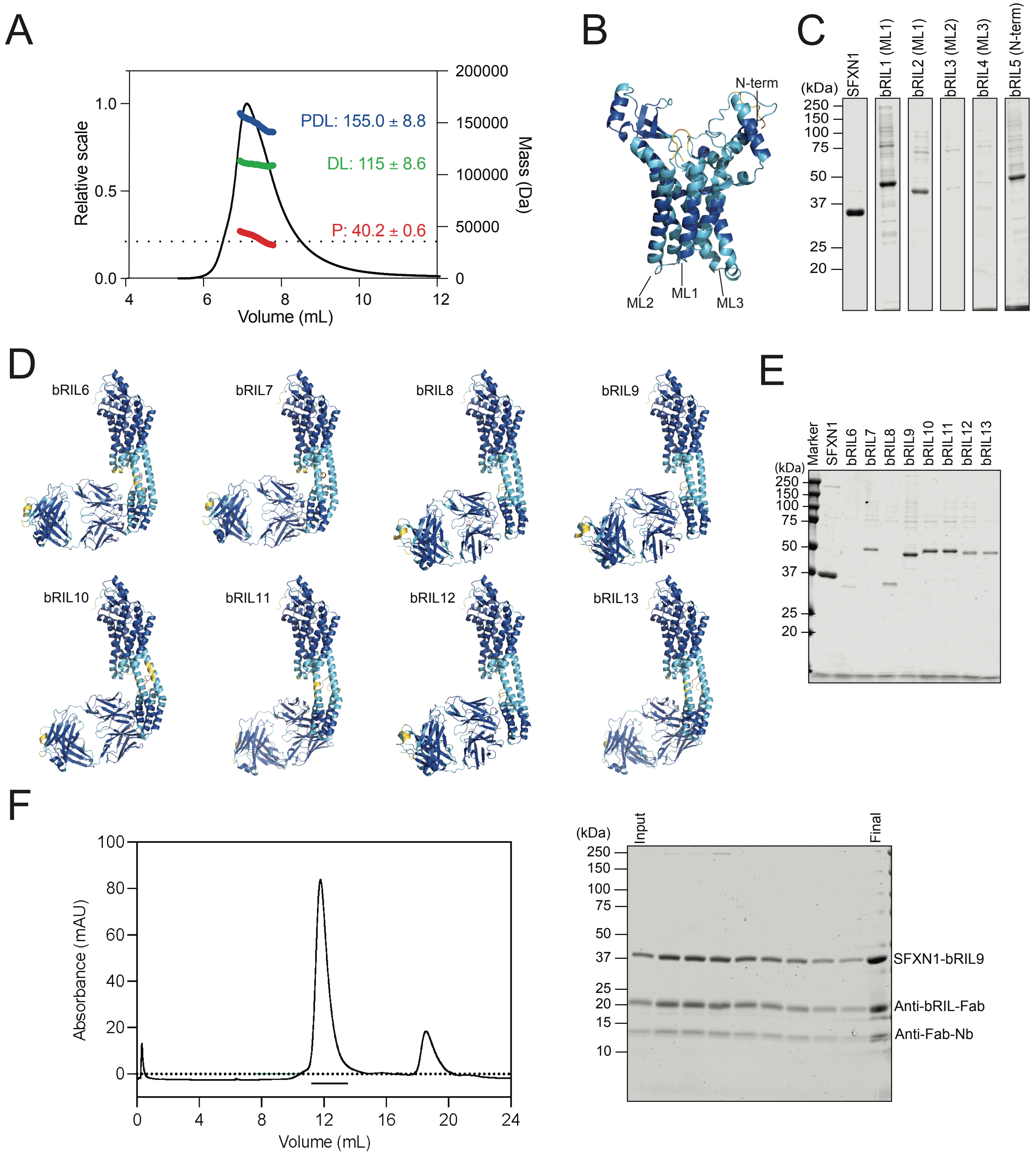


**Supplementary Figure 1: SFXN1 biochemical characterisation and bRIL fusion engineering by protein design for structural studies.** (**A**) SEC-MALLS analysis of SFXN1 showing the mass of the protein (P; red), the associated protein-detergent-lipid (PDL; blue) and detergent-lipid alone (DL; green). A representative chromatogram is shown. Numbers indicate the mean mass (kDa) from three independent experiments (n=3) and the standard deviation. (**B**) AlphaFold2 prediction of SFXN1, bRIL insertion sites in matrix loops 1-3 (ML1-3) and at the N-terminus are shown (left). (**C**) small scale purification of wild-type SFXN1 and bRIL fusion constructs (bRIL1-5). (**D**) AlphaFold2 models of designed SFXN1 BRIL6-13 fusions inserted in ML1. (**E**) SDS-PAGE analysis of purified wild-type SFXN1 and bRIL6-13 fusion constructs. (**F**) Size-exclusion chromatography (SEC) profile and corresponding SDS–PAGE analysis confirming the purity and monodispersity of the SFXN1–bRIL9:anti-bRIL Fab:anti-Fab nanobody complex. AF2 models are coloured by predicted local distance difference test (pLDDT) confidence scores (dark blue for 90-100 and very high confidence; light blue for 70-90 and confident, yellow for 50-70 and low confidence and orange for <50 and very low confidence).

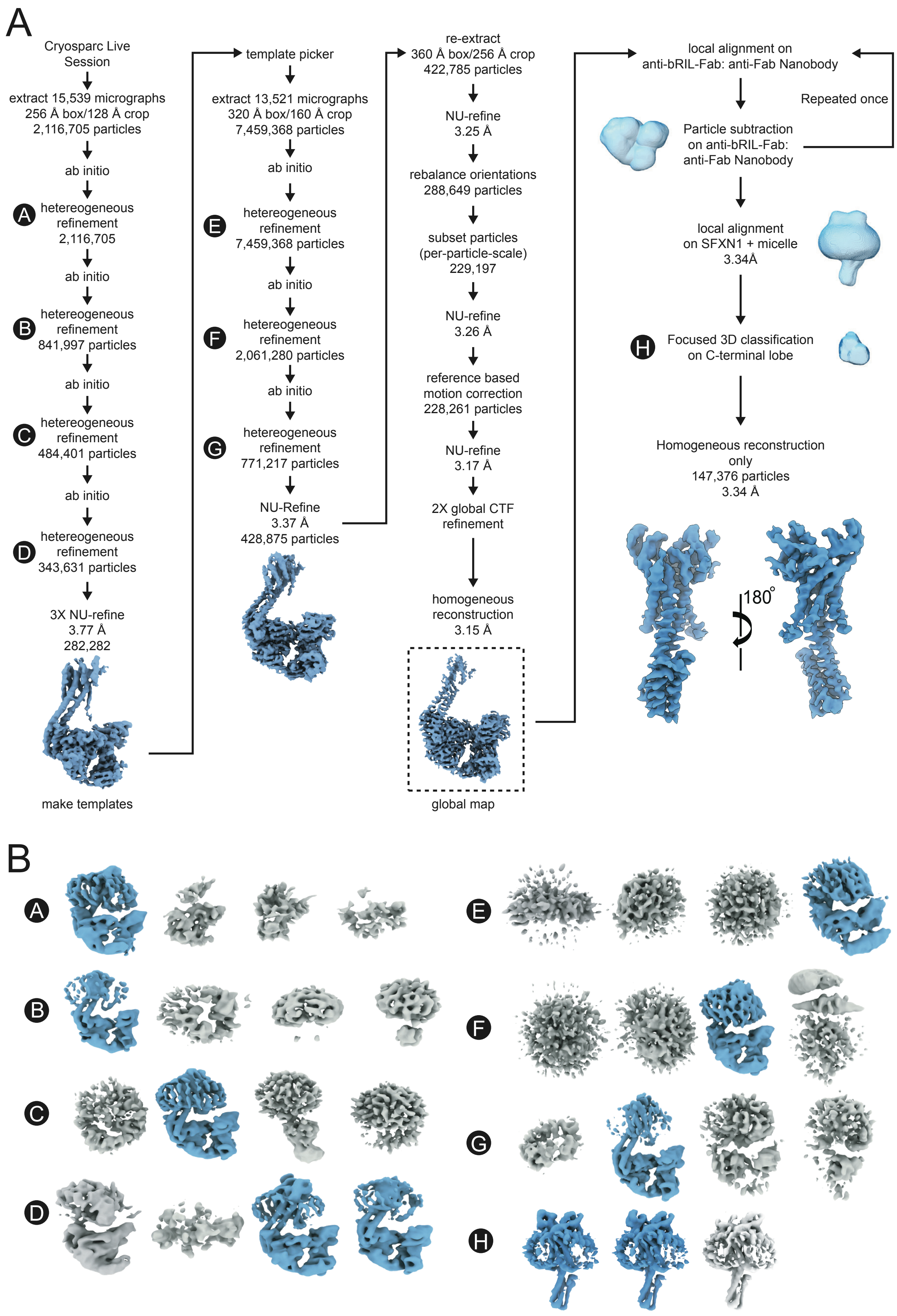


**Supplementary Figure 2: Cryo-EM data processing workflow.** (**A**) CryoSPARC processing workflow used to obtain the global map of the SFXN1–bRIL9:anti-bRIL Fab:anti-Fab nanobody complex and the locally refined SFXN1-bRIL9 map. The number of micrographs, particle counts, box size, Fourier resampling parameters, and resolutions (reported at GS-FSC = 0.143) are indicated at each stage. Key intermediate maps are shown as blue surface renderings. Heterogeneous refinements are labelled A–G (black circles with white text), and 3D classification without alignment as H. Masks used during refinement are shown as light blue surfaces. (**B**) Heterogeneous refinement classes (A–G) and focused 3D classification (H) results. Classes selected for further analysis are shown in blue, and discarded classes in grey, along with their associated particle counts.

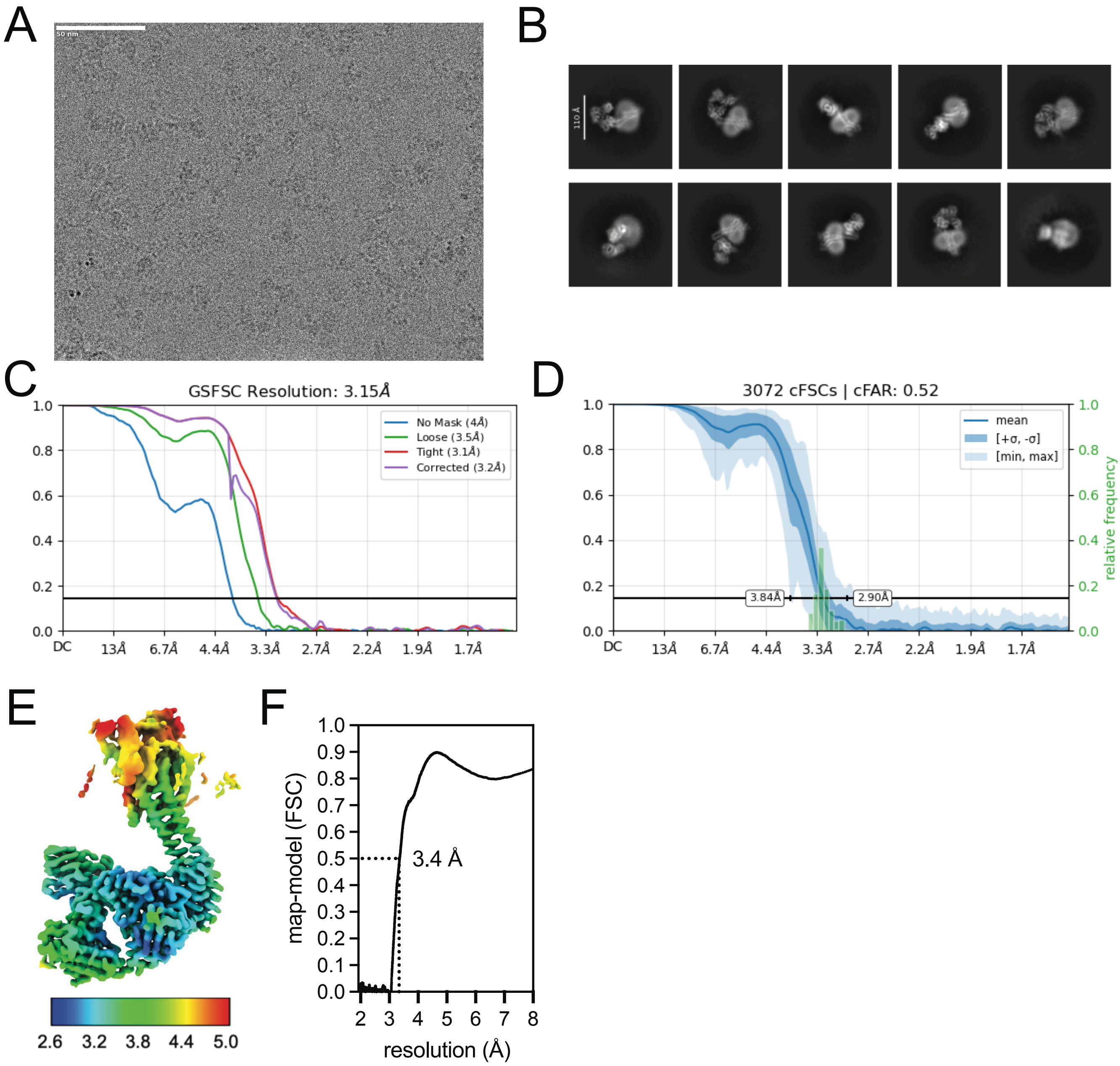


**Supplementary Figure 3: Map quality for Sideroflexin 1-bRIL9:anti-bRIL Fab:anti-Fab nanobody complex (SFXN1 complex).** (**A**) Example micrograph from data collection on Titas Krios. (**B**) Selected 2D class averages from the particles used to resolve the final SFXN1 complex reconstruction. (**C**) GS-FSC calculated in cryosparc for SFXN1 complex map. (**D**) A directional cFSC (conical FSC) calculated in cryosparc for SFXN1 complex map. (**E**) A local resolution surface rendering of the SFXN1 complex; local resolution is rendered as rainbow colouring from 2.6 Å to 5.0 Å on a sliding scale from blue to red. (**F**) Map-model FSC calculated for SFXN1 complex with a protein model including SFXN1 residues (105-109) from TM1 and (116-120) from TM2 connecting to bRIL in complex with the anti-bRIL Fab and anti-Fab nanobody. Dashed line indicates the corresponding resolution at FSC = 0.5.

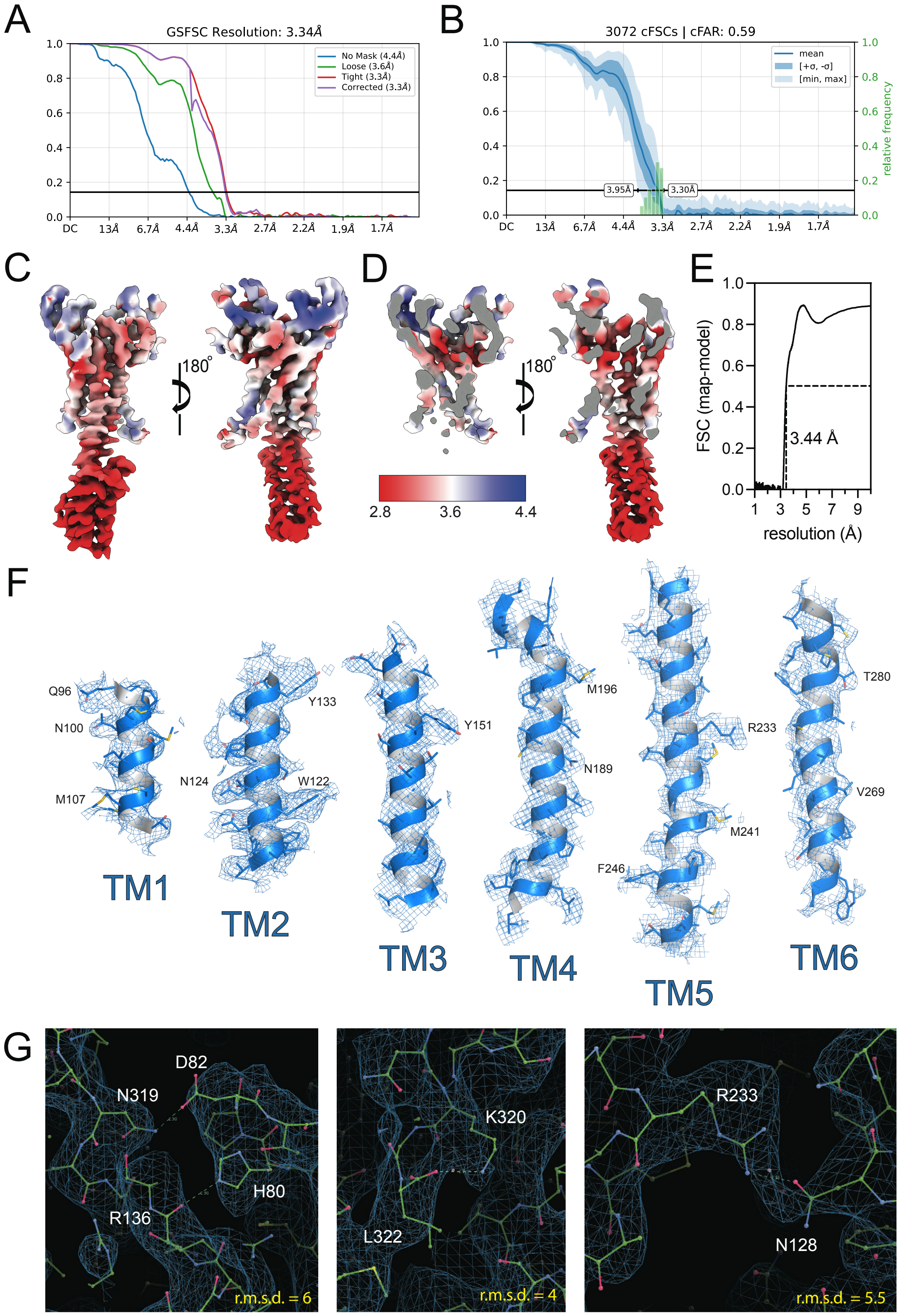


**Supplementary Figure 4: Cryo-EM map quality after particle subtraction and local refinement on SFXN1-bRIL9** (**A**) GS-FSC calculated by CryoSPARC for SFXN1-bRIL9 map. (**B**) A directional cFSC (conical FSC) calculated by CryoSPARC for SFXN1-bRIL9 map. (**C**) A local resolution surface rendering of SFXN1 complex. Local resolution is shown as a sliding scale from red to white to blue on a sliding scale from 2.8 Å to 4.4 Å, and (**D**) a mid-protein slice. (**E**) map-model FSC calculated for SFXN1-bRIL9, dashed line indicates resolution at FSC = 0.5. (**F**) Local map-model fits of transmembrane helices. SFXN1 TMs are shown as blue cartoons and sticks with CPK colouring. The cryosparc sharpened map is shown at 5 r.m.s.d. (**G**) Local map fit of the HPDT motif (left), K320 to L322 (center), N128 to R233 (right) interactions shown in coot. Residues are shown as green sticks with CPK colouring, the map is shown as a blue mesh and dashes indicated proposed bonds. The map r.m.s.d is shown as an inset.

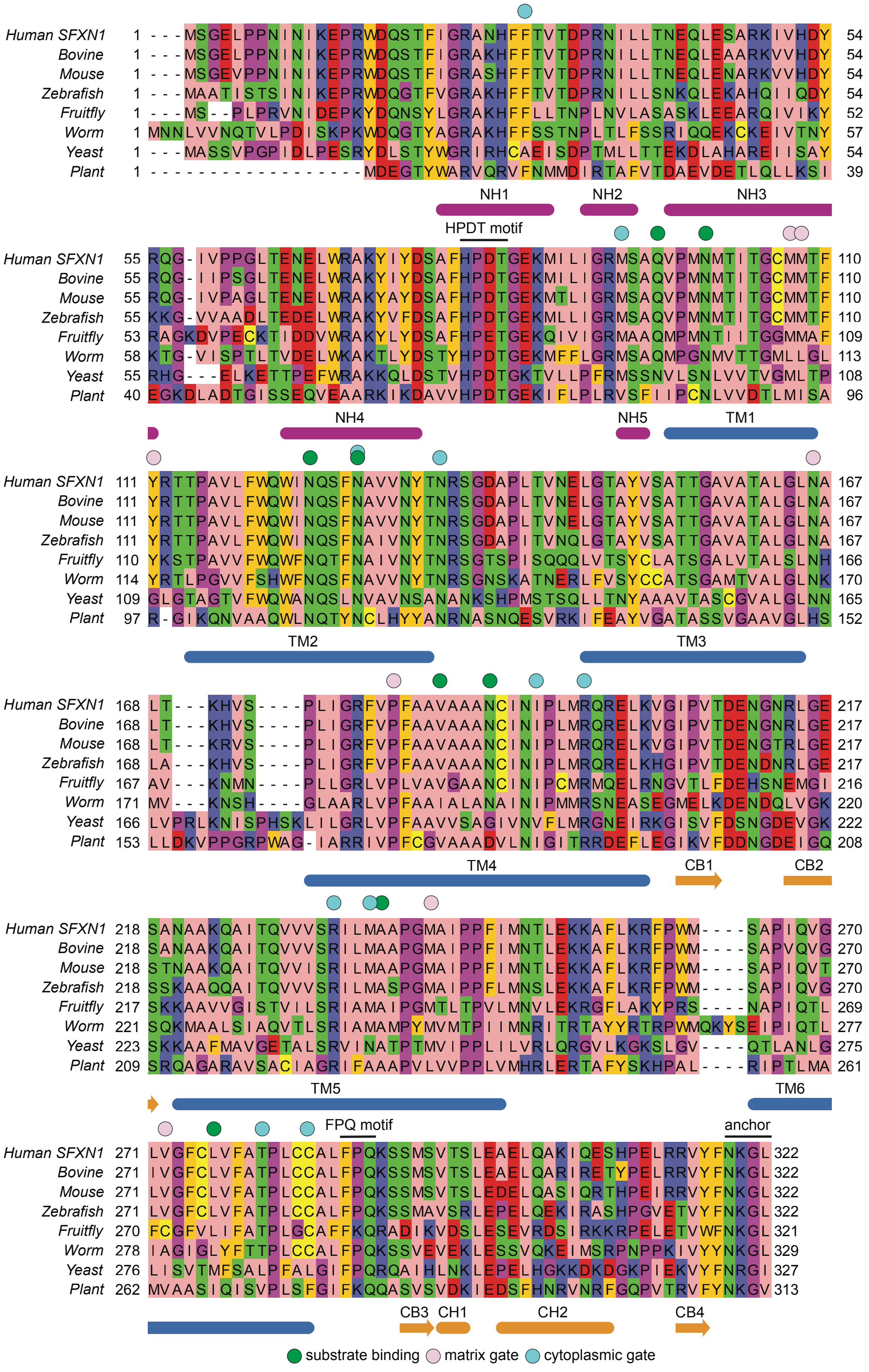


**Supplementary Figure 5: SFXN1 focused alignment across species orthologues.** *Homo sapiens* (Human SFXN1), *Bos taurus* (Bovine), *Mus musculus* (Mouse), *Danio rerio* (Zebrafish), *Drosophila melanogaster* (Fly), *Caenorhabditis elegans* (Worm), *Saccharomyces cerevisiae* (Yeast) and *Physcomitrium patens* (Plant) protein sequences were retrieved from Uniprot, aligned with MUSCLE and manually curated in Jalview. Residues are coloured according to the zappo colouring scheme: aliphatic are pink: hydrophillic are green; aromatic are orange; basic are blue; acidic are red; proline and glycine are purple; cysteine is yellow. Secondary structure is annotated below the alignment as a rounded rectangle for helices and arrows for beta-sheets, and coloured according to Figure 1. The HPDT, FPQ and anchor motifs are identified with a solid black line and text above the alignment. A legend is displayed at the foot of the figure to illustrate the coloured circles for structural elements described. Substrate binding site residues are green; matrix gate residues are pink; cytoplasmic gate residues are blue.

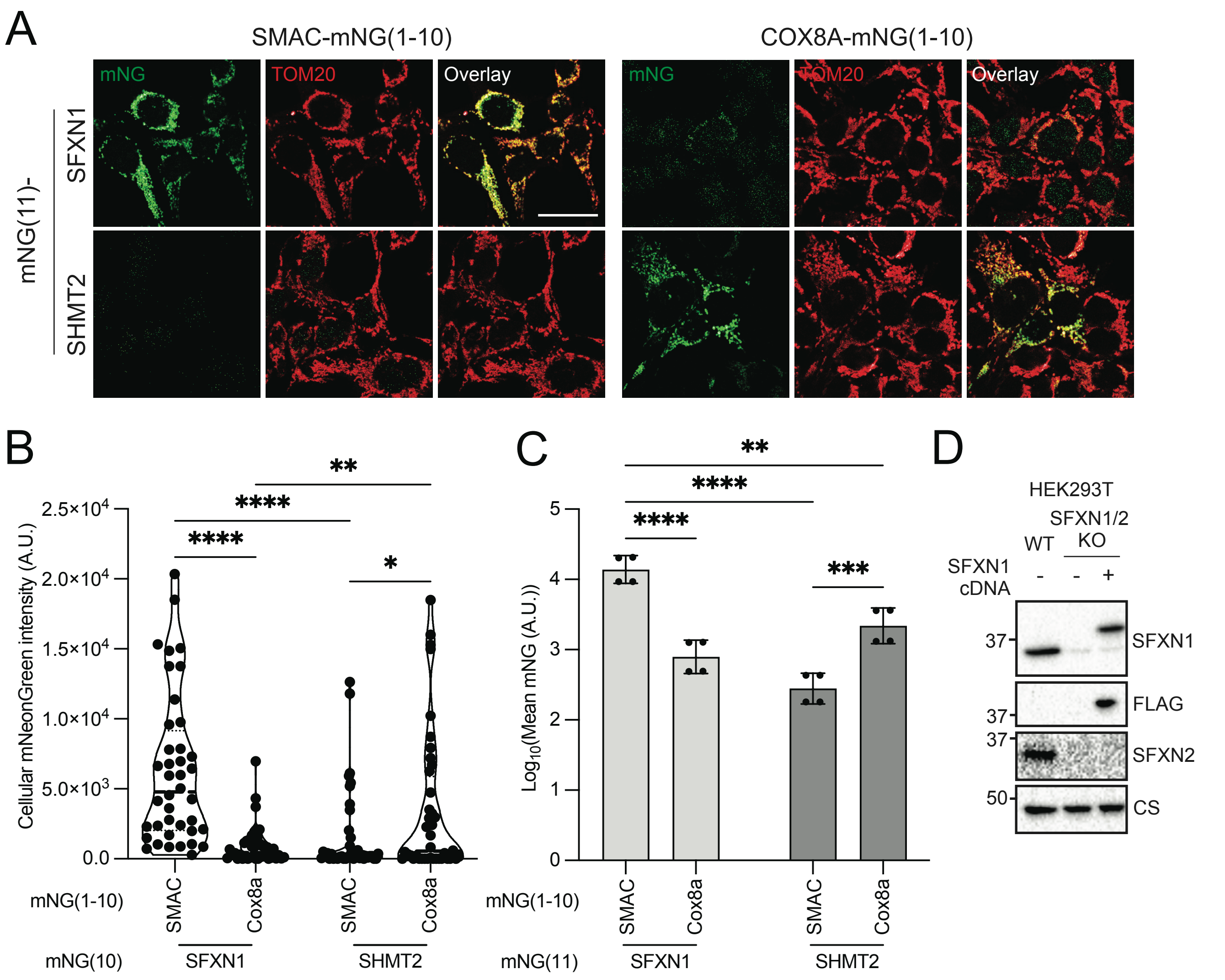


**Supplementary Fig. 6: Topology of SFXN1 in mitochondria and engineering of HEK293T SFXN1/2 knockout cell lines.** (**A**) Confocal microscopy images of cells stably expressing mNG(11)-SFXN1 or SHMT2-mNG(11) fusions, co-expressed with SMAC-mNG(1-10) or Cox8A-mNG(1-10) fusions. TOM20 staining marks mitochondria. Scale bar, 10 µm. (**B**) Quantification of fluorescence signal shown as violin plots, with individual cells (n=40) indicated as dots; center lines denote the median and quartiles. (**C**) Flow cytometry analysis of HEK293T cells shown in (b) (n=4 independent experiments; 50,000 cells per experiment). (**D**) Validation of CRISPR-Cas9-edited HEK293T cell lines. SFXN1 and SFXN2 knockout cells were generated as described in the Methods. Immunoblot analysis was performed using antibodies against SFXN1, SFXN2, FLAG and citrate synthase (loading control). For (b,c), statistical significance was assessed by two-way analysis of variance (ANOVA) with Šídák's multiple-comparison post hoc test (*P < 0.05; **P < 0.01;***P < 0.001;****P < 0.0001).

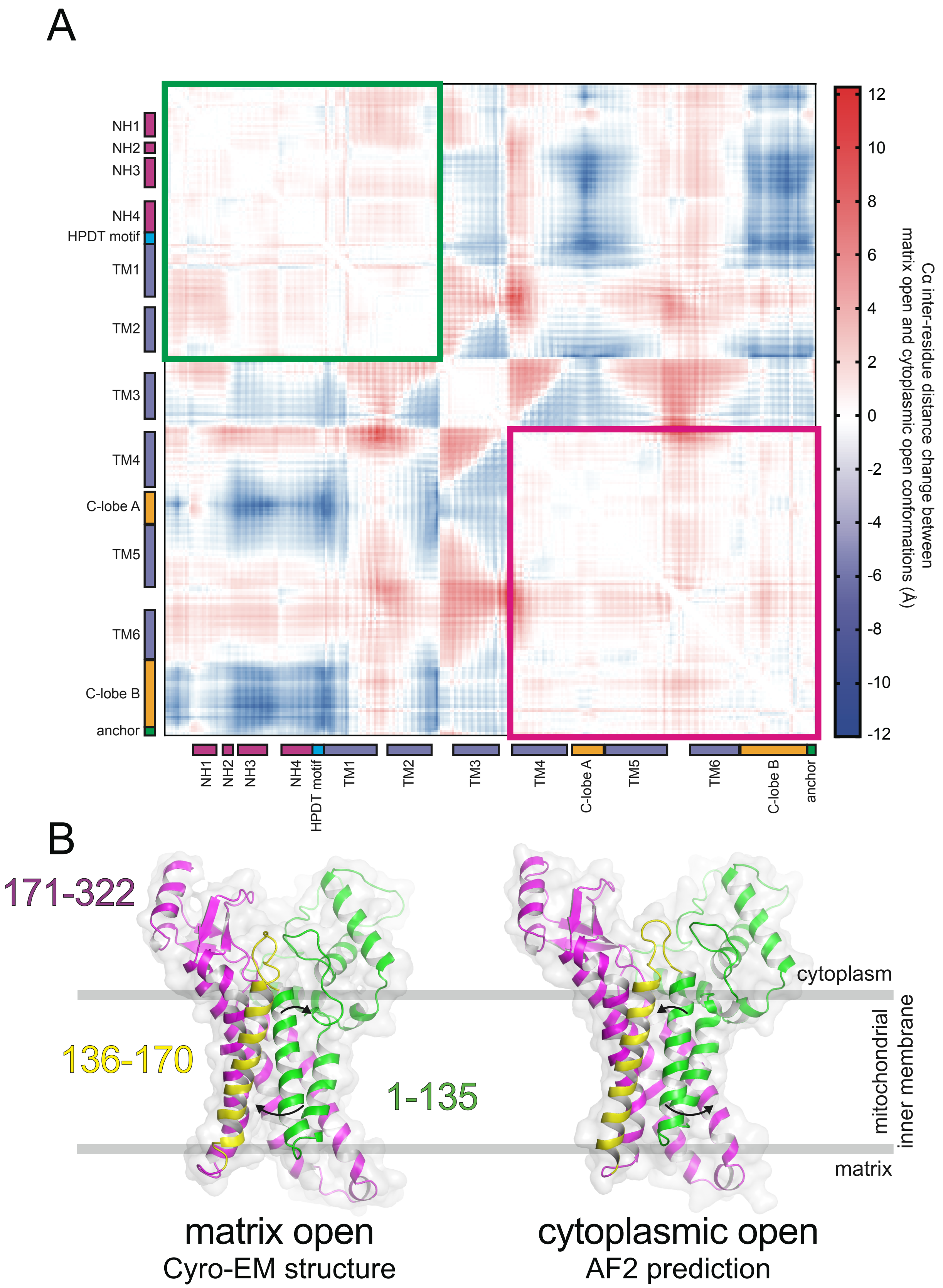


**Supplementary Figure 7: Motions of SFXN1 between cytoplasmic open and matrix open states.** (**A**) Difference contact map between the matrix open and cytoplasmic open state. We calculated the intra-residue Cα between all residues for each conformation and then calculated the difference between conformations (matrix open intra-residue difference minus cytoplasmic open intra-residue difference). The differences are displayed as a heatmap coloured on a blue to red sliding scale from -12 Å to + 12 Å. Elements from SFXN1 are shown and coloured as in Fig. 1. Green square indicates the rigid body formed by residues 1-135, and magenta square indicates residues 171-322. (**B**) SFXN1 in the cytoplasmic-open and matrix-open conformation was globally aligned and coloured green for residues 1-135, yellow for 136-170 and magenta for 171-322. Arrows indicate movement between conformations of residues 1-135.

**Table S1, related to Figure 1. Cryo-EM data collection, refinement and validation statistics.**

| | |
|---|---|
| **Human Sideroflexin 1**<br>EMD-73540 ([https://www.ebi.ac.uk/emdb/EMD73540).<br>PDB 9YW8 ([http://doi.org/9YW8]) |  |
| **Data collection and processing** | |
| Voltage (kV) | 300 |
| Electron exposure ($e^-$/Å$^{-2}$) | 50.93 |
| Defocus range (µm) | 0.8 – 2.0 |
| Pixel size (Å) | 0.825 |
| Symmetry imposed | C1 |
| Initial particle images (no.) | 7,459,368 |
| Final particle images (no.) | 147,376 |
| FSC threshold | 0.143 |
| **Refinement** | |
| Initial model used | Alphafold2 SFXN1-bRIL9 (local run) |
| Map Resolution (Å) | 3.34 |
| Map Resolution range (Å)<br>(per atom position) | 2.40-4.40 |
| **Model composition** | |
| Non-hydrogen atoms | 3,272 |
| Protein residues | 426 |
| **B factors (Å$^2$)** | |
| Protein | 247 |
| **R.M.S.D. deviations** | |
| Bond lengths (Å) | 0.013 (0) |
| Bond angles (°) | 1.733 (0) |
| **Validation** | |
| MolProbity score | 0.81 |
| Clashscore | 1.07 |
| Poor rotamers (%) | 0.89 |
| CaBLAM outliers (%) | 0.96 |
| CCmask | 0.84 |
| CCvolume | 0.84 |
| Map-to-model FSC (FSC = 0.5) | 3.45 |
| Q-score | 0.47 |
| **Ramachandran plot** | |
| Favoured (%) | 98.34 |
| Allowed (%) | 1.42 |
| Disallowed (%) | 0.24 |
